# Strategic stabilization of arousal boosts sustained attention

**DOI:** 10.1101/2022.03.04.482962

**Authors:** J.W. de Gee, Z. Mridha, M Hudson, Y. Shi, H. Ramsaywak, S. Smith, N. Karediya, M. Thompson, K. Jaspe, H. Jiang, W. Zhang, M. J. McGinley

## Abstract

Changes in autonomic arousal, such as mounting sleep pressure, and changes in motivation, such as fluctuating environmental reward statistics, both profoundly influence behavior. Our experience tells us that we have some capacity to control our arousal when doing so is important, such as staying awake while driving a motor vehicle. However, little is known about how decision computations are jointly influenced by arousal and motivation, including whether animals, such as rodents, can adapt their arousal state to their needs. Here, we developed and show results from an auditory feature-based sustained-attention task with intermittently shifting task utility. We use pupil size to estimate arousal across a wide range of states and apply novel signal detection theoretic and accumulation-to-bound modeling approaches in a large behavioral cohort. We find that both pupil-linked arousal and task utility have major impacts on multiple aspects of performance. Although substantial arousal fluctuations persist across utility conditions, mice partially stabilize their arousal near an intermediate, and optimal, level when task utility is high. Behavioral analyses show that multiple elements of behavior improve during high task utility and that arousal influences some, but not all, of them. Specifically, arousal influences the likelihood and timescale of sensory evidence accumulation, but not the quantity of evidence accumulated per time step while attending. In sum, the results establish specific decision-computational signatures of arousal, motivation, and their interaction, in attention. So doing, we provide an experimental and analysis framework for studying arousal self-regulation in neurotypical brains and diseases such as attention-deficit/hyperactivity disorder.

## Introduction

Arousal state and motivational state both profoundly impact behavior, but how they interact at the level of decision computations is largely unknown. Autonomic arousal, controlled by neuromodulators released from the reticular activating system, fluctuates on multiple time scales, shaping sensory processing and decision-making behavior (Aston-Jones & Cohen, 2005; Harris & Thiele, 2011; Lee & Dan, 2012; McGinley, Vinck, et al., 2015). Fluctuations in pupil size at constant luminance track global arousal state (Joshi & Gold, 2020; McGinley, Vinck, et al., 2015) and the activity of the underlying neuromodulatory systems, including noradrenaline (Breton-Provencher & Sur, 2019; de Gee et al., 2017; Joshi et al., 2016; Murphy et al., 2014; Reimer et al., 2016; Varazzani et al., 2015) and acetylcholine (de Gee et al., 2017; Mridha et al., 2021; Reimer et al., 2016). Optimal behavioral and neural detection of simple sounds occurs at intermediate levels of spontaneously fluctuating pupil-linked arousal (Beerendonk et al., 2023; McGinley, David, et al., 2015; Schriver et al., 2018). Large baseline pupil size is associated with task disengagement, including exploratory behaviors like locomoting (Gilzenrat et al., 2010; Jepma & Nieuwenhuis, 2011; McGinley, David, et al., 2015) and small baseline pupil is associated with drowsiness or sleep (McGinley, David, et al., 2015; Yüzgeç et al., 2018). This ‘inverted-U’ shape of the effects of pupil-linked arousal on task exploitation, versus task disengagement as exploration or rest, is referred to as the Yerkes-Dodson relationship and relates closely to the ‘adaptive gain’ theory of noradrenergic tone (Arnsten & Li, 2005; Aston-Jones & Cohen, 2005; Berridge & Waterhouse, 2003; Kane et al., 2017; Pfeffer et al., 2021; Usher et al., 1999; Yerkes & Dodson, 1908).

Sustained and feature-based attention is an important cognitive function that is potently influenced by motivational state and can be effectively modeled in mice (Robbins, 2002; L. Wang & Krauzlis, 2018). Motivation level, for example driven by the magnitude of reward for attending effectively, has been shown to impact the intensity aspect of attention, also called ‘attentional effort’ (Ghosh & Maunsell, 2021; Sarter et al., 2006). In lay terms, motivation level can turn attention level up and down. We hereafter refer to this psychological construct as ‘attentional intensity’. Signal detection theory (SDT) analysis can capture motivated shifts in attentional intensity as changes in the discrimination sensitivity, or d’, in contrast to orthogonal changes in behavioral strategy, which are captured as changes in bias, or c (also called criterion) (Green & Swets, 1966). In attention tasks, as with most decision-making processes, effects on d’ and bias are thought to arise from changes in the underlying process of accumulating decision-relevant sensory evidence, implemented in a distributed network of brain areas (Shadlen & Kiani, 2013; Steinmetz et al., 2019; van Vugt et al., 2018). In widely used accumulation-to-bound models of judgments about weak time-varying sensory signals in noise, sensory cortex encodes the noise-corrupted decision evidence, and downstream association and motor cortices accumulate this noisy sensory signal into a decision variable that determines the behavioral choice (Bogacz et al., 2006; Shadlen & Kiani, 2013; Siegel et al., 2011; X.-J. Wang, 2008). Thus, SDT and accumulation-to-bound models can provide basic and high-resolution accounts, respectively, of attention-related decision-making processes.

How motivated attention interacts with this inverted-U model of pupil-linked neuromodulatory system function is not known. This is an important gap in knowledge, because our experience tells us that motivated self-control of arousal importantly impacts attention. For example, dysregulated self-control of arousal is a prominent aspect of attention-deficit/hyperactivity disorder (ADHD) and other developmental disabilities and psychiatric disorders (de Lecea et al., 2012; Sander et al., 2015; Zhao et al., 2022). Thus, a central and unsolved question about the role of arousal in brain function is how spontaneous arousal fluctuations, driven by internal metabolic and/or memory consolidation demands (Squire et al., 2015), are strategically influenced to match arousal state to task demands (Aston-Jones & Cohen, 2005). According to the adaptive gain theory, when task utility (i.e. reward expectation) is high, one should regulate towards a moderate arousal state to facilitate optimal task engagement. In contrast, when utility is low, one should upregulate to a high arousal state to facilitate exploration of alternatives, or downregulate to a low arousal state to rest and consolidate. In line with this proposed adaptability of arousal control, the locus coeruleus receives strong projections from frontal regions including the orbital frontal cortex and anterior cingulate cortex (Arnsten & Goldman-Rakic, 1984; Aston-Jones & Cohen, 2005; de Gee et al., 2017; Joshi & Gold, 2022; Porrino & Goldman-Rakic, 1982) and has more complex organization than previously assumed (Breton-Provencher & Sur, 2019; Poe et al., 2020; Totah et al., 2018). These organizational features may provide a neural substrate for strategic self-regulation of arousal.

We here sought to understand how fluctuations in arousal and shifts in task utility interact to influence decision computations supporting sustained feature-based attention. We developed a ‘sustained-attention value’ task for head-fixed mice and report behavioral and pupillary signatures of in a large cohort. The task manipulates coherent acoustic motion in time-frequency space, analogous to random-dot motion used extensively in the visual system (Newsome et al., 1989). We develop and apply a survival analysis-based SDT approach, as well as tailored accumulation-to-bound models, and assess the interacting effects of task utility and pupil-linked arousal. We find that both arousal state and motivational state have large impacts on decision computations. During periods of high task utility mice exhibit multiple signatures of heightened attentional intensity, which are partially mediated by stabilization of arousal closer to an optimal, moderate levels.

## Results

### A feature-based sustained-attention task with nonstationary task utility

To study how arousal and motivation interact to shape decision computations during sustained attention, we developed a quasi-continuous auditory detection task for head-fixed mice (**Fig. 1**). Mice were trained to detect coherent time-frequency motion (the ‘signal’; called temporal coherence (Shamma et al., 2011)) embedded at unpredictable times in an ongoing and otherwise random cloud of tones (the ‘noise’). The task required sustained, attentive listening to achieve high detection performance, due to the perceptual difficulty of noticing the temporally unpredictable emergence of a high-order acoustic feature. Mice were motivated to elicit lick responses by being food scheduled and by administration of sugar-water reward when they licked during the signal. To suppress excessive licking, mice received a 14-second timeout if they licked during the noise (**Fig 1A**). We manipulated the utility of performing the task by shifting the reward size back and forth between 60 trials of high (12 µl) and 60 trials of low (2 µl) values, in 7 consecutive blocks. We recorded the timing of correct (‘Hit’) and incorrect (‘False Alarm’) responses with respect to the ongoing sound stimuli (**Fig 1B**, top rows). We also measured the diameter of the pupil as readout of arousal and walking speed as an additional behavioral state measure (**Fig 1B**, bottom rows). Hereafter, we refer to the task as the sustained-attention value task (see Methods for additional details).

**Figure 1.**
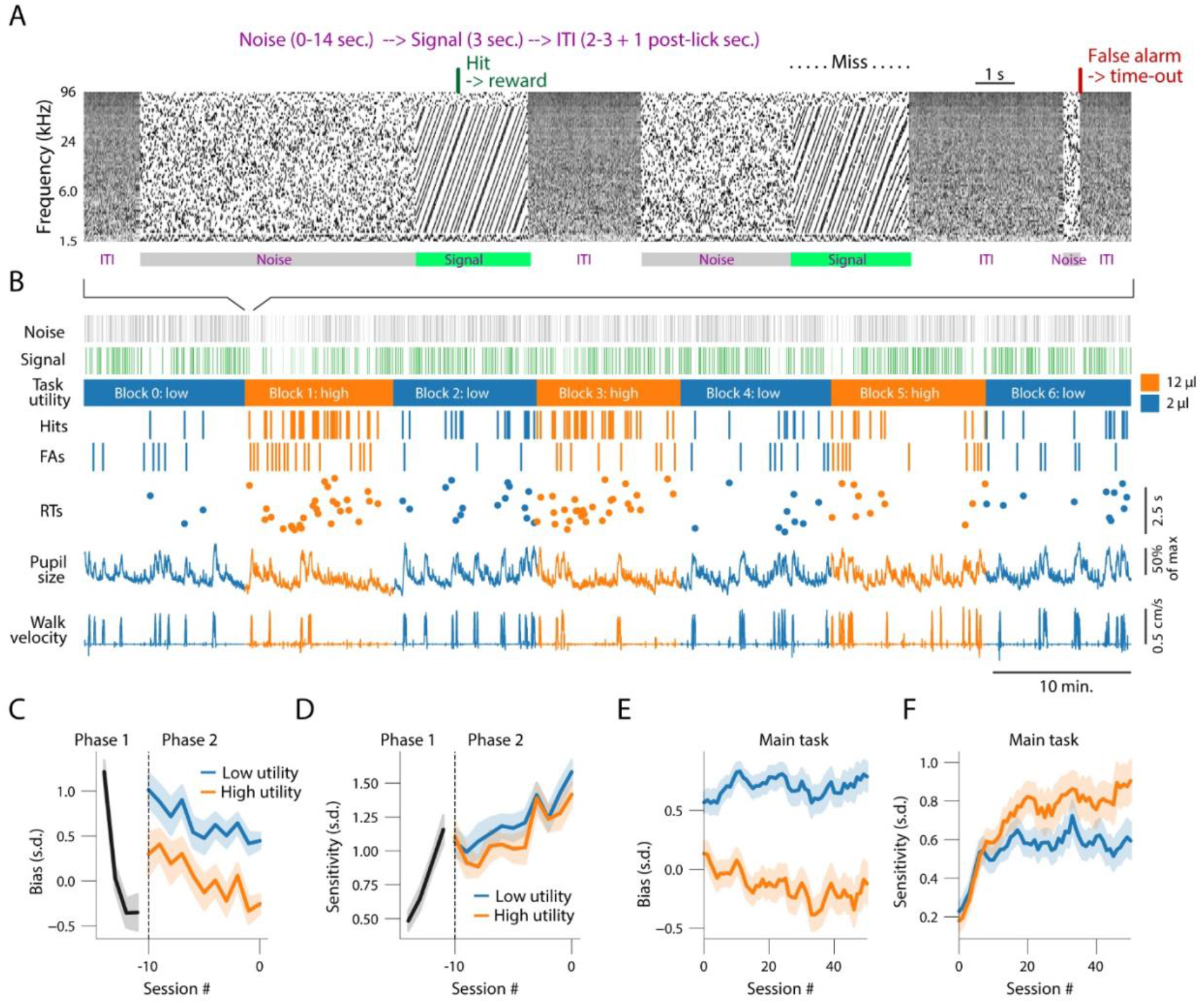
Monitoring performance and pupil-linked arousal during a sustained-attention value task. **(A)** Spectrogram of the sound during an example of three consecutive trials. Correct go responses (hits, green text and arrow at top) were followed by 2 or 12 μL of sugar water. Reward magnitude alternated between ‘low’ and ‘high’ values (2 and 12 μL, respectively) in blocks of 60 trials. Incorrect go-responses (false alarms, red text and arrow at top) terminated the trial and were followed by a 14 s timeout, indicated to the animal as pink acoustic noise. **(B)** Example session. From top to bottom: noise stimulus timing, signal stimulus timing, reward/utility context, correct responses (hits), incorrect responses (false alarms), reaction times (RTs) on hit trials, pupil size and walking velocity. Color (blue or orange) indicates utility/reward context. **(C)** Bias (Methods) across experimental sessions in learning phases 1 and 2 (Methods); session numbers are with respect to the last session in phase 2. **(D)** As C, but for sensitivity (Methods). **(E,F)** As C,D, but for the main task (Methods). See Fig. S1 for fitted learning curves and time constants. Panels C-F: shading, 68% confidence interval across animals (N=88, n=4473 sessions).

To assess the basic patterns of how mice learned and performed the sustained-attention value task, we applied signal detection theory to assess perceptual sensitivity (d’, also called discriminability) and bias (c, also called criterion). Because the duration of the noise varied between trials, and because each signal was preceded by a noise stimulus on each trial, care was required in calculating hit and false alarm rates. Hit rate (HR) was calculated in the traditional manner, as the fraction of signals for which the mouse licked. Because signals only occurred if the mouse did not lick during the noise, this HR is implicitly a conditional probability. The appropriate false alarm rate (FAR) was calculated as the time-in-noise matched conditional probability, but without signal present, which could be estimated using the Kaplan-Meier Survival function (see Methods for details). Sensitivity and bias were defined as the difference or sum, respectively, of the z-scored HR and FAR (Green & Swets, 1966). We also measured reaction time, as the latency from signal start to first lick on hit trials, and reward probability, as the fraction of trials that ended in reward.

Because the sustained-attention value task is, by design, demanding of attentional resources, we hypothesized that mice would increase their attentional intensity during blocks of high reward, to exploit the high utility, and reduce attentional intensity in low-reward blocks, to conserve cognitive resources and/or engage in other activities. To test whether mice could learn to adaptively adjust their attentional intensity, we measured sensitivity and bias across learning of the task. Training occurred in three phases. In phase 1, the signal was 6 dB louder than the noise, several free-reward trials (not contingent on response) were administered, and reward magnitude was constant (5 μL) throughout the session. During this phase, mice learned to respond (lick) to harvest rewards; within just three sessions their bias changed from quite conservative (positive bias values) to slightly liberal (negative bias values; **Fig. 1C**). Sensitivity also increased across the first few sessions (**Fig. 1D**). In phase 2, we introduced the block-based shifts in reward magnitude and trained the mice until they reached a performance threshold (Methods). Sensitivity gradually increased during phase 2 (**Fig. 1D**).

In phase 3, which was the main task, acoustic signal and noise stimuli were of equal loudness and the small number of classical conditioning trials were removed. By this time, mice were sufficiently experienced with the task to maintain engagement (**Fig. 1E**). However, since they could no longer rely on the small loudness difference between signal and noise sounds, their sensitivity dropped to near zero at the beginning of phase 3 (**Fig. 1F**). Re-learning the, time-varying and now purely feature-based, signal detection was perceptually difficult, as intended, as indicated by the shallow learning curves for sensitivity and reward probability (**Fig. 1F**; **Fig. S1G,K**). Thus, mice rapidly learned the task structure, but slowly learned the feature-based attention and adaptive attentional intensity allocation (**Fig. 1C**-**F**; **Fig. S1E-L**; see Methods for details).

Due to the temporal structure of the task, it is possible that mice could learn to match their lick times to the temporal statistics of signal occurrence to harvest at least some reward, rather than attend to the temporal coherence per se. To rule out such a timing-based strategy, and demonstrate feature-based listening, we trained a cohort of mice (N=10; n=142 sessions) in a psychometric variant of the task in which signal coherence was varied randomly from trial to trial, in addition to the block-based utility manipulation. Signal coherence was degraded by manipulating the percentage of tones in each chord that moved coherently through time-frequency space (**Fig. S1M**; Methods). This manipulation is analogous to reducing spatial motion coherence in visual random-dot motion tasks (Newsome et al., 1989). As would be expected if the mice were employing feature-based attention, rather than a timing-based strategy, both sensitivity and reward probability lawfully varied with signal coherence (**Fig. S1N,O**). Further supporting feature-based attention, rather than a temporal strategy, empirical sensitivity (Fig. 1E,F) and reward probabilities (**Fig. S1J, top**) were substantially higher than those predicted with random licking (**Fig. S2A**) and signal start-time sorted survival functions for hits deviated strongly from the time-matched false-alarm survival functions (**Fig. S2B**).

### Upregulation of task engagement and attentional intensity during high task utility

We next analyzed if mice strategically regulate attentional intensity across blocks. We interpreted block-based shifts in sensitivity as an indication of changing attentional intensity and block-based shifts in bias as changing task engagement. Therefore, we measured the time scales (both within and across blocks) on which mice changed their bias and sensitivity to shifts in task utility (Fig. 2A-F). We observed that mice spent most of the 1^st^ block (termed block ‘0’; always low reward) becoming gradually engaged in the task (Fig. 2A,D,G,J; see also Fig. 5A). Therefore, we focused all analyses on the subsequent six blocks (termed blocks ‘1’ through ‘6’), for which the correct lick-triggered reward size alternated in blocks between 12 μL (high reward, orange traces) and 2 μL (low reward, blue traces; see Methods for additional details).

**Figure 2.**
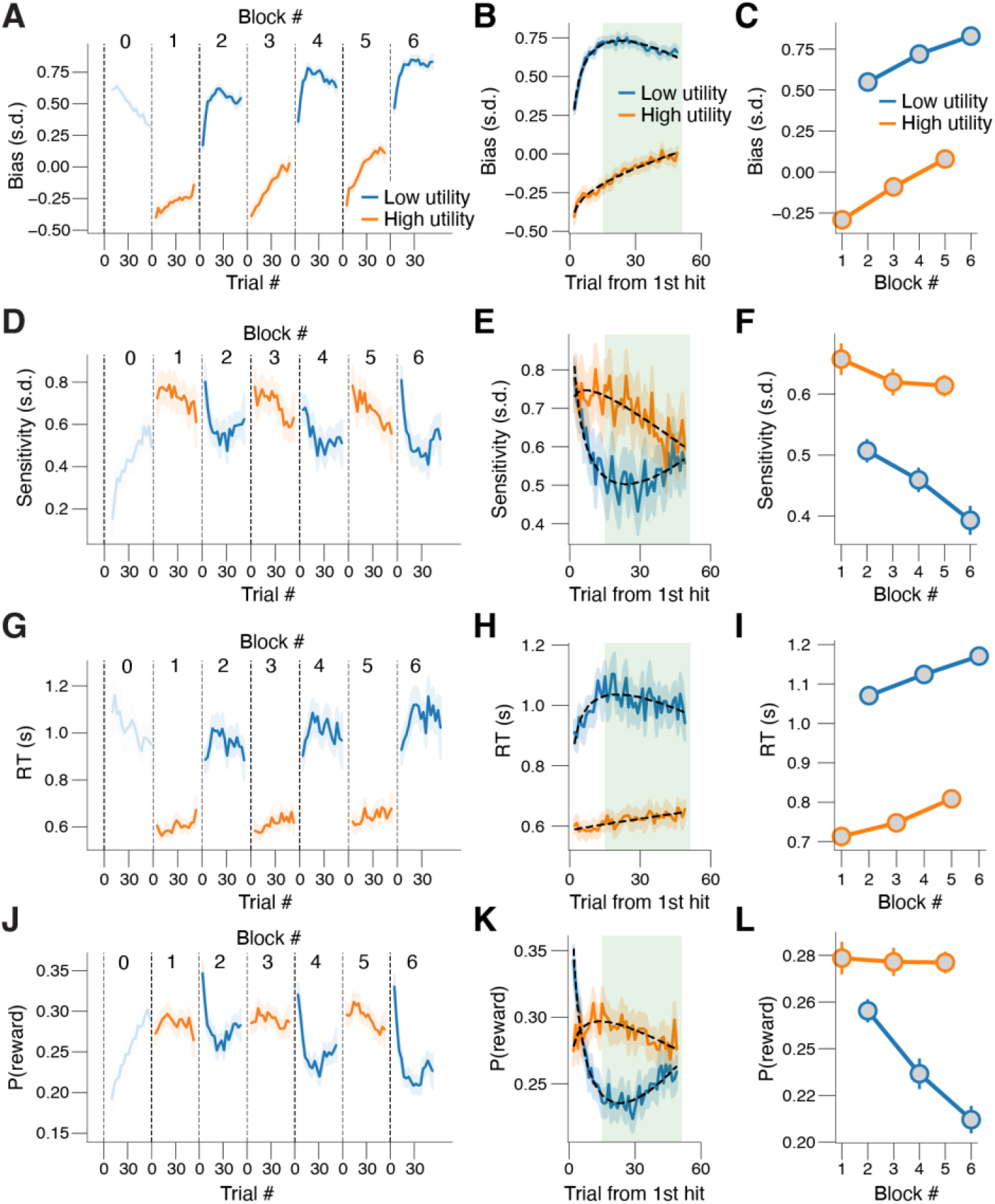
Rapid and adaptive changes in performance after shifts in task utility. **(A)** Bias across low-reward and high-reward blocks in each experimental session, locked to first hit in block. Data from the first block of each session (low utility; termed block ‘0’) was excluded from all analyses, as mice spent this block becoming engaged in the task (see also panels D,G and J; Methods). **(B)** As A, but collapsed across blocks of same reward magnitude. The green shaded area indicates the trials used when pooling data across trials within a block (e.g. panel C). **(C)** As A, but collapsed across trials within a block. Stats, 2-way repeated measures ANOVA (factors task utility [high vs. low] and time-on-task [early, middle, late]); main effect task utility: F_1,87_ = 657.2, p < 0.001; main effect time-on-task: F_2,174_ = 124.5, p < 0.001; interaction effect: F_2,174_ = 4.0, p = 0.020. **(D-F)** As A-C, but for sensitivity. Main effect task utility: F_1,87_ = 33.2, p < 0.001; main effect time-on-task: F_2,174_ = 7.8, p = 0.001; interaction effect: F_2,174_ = 2.6, p = 0.076. **(G-I)** As A-C, but for RT. Main effect task utility: F_1,87_ = 499.3, p < 0.001; main effect time-on-task: F_2,174_ = 27.4, p < 0.001; interaction effect: F_2,174_ = 0.3, p = 0.741. **(J-L)** As A-C, but for reward probability. Main effect task utility: F_1,87_ = 19.7, p < 0.001; main effect time-on-task: F_2,174_ = 26.0, p < 0.001; interaction effect: F_2,174_ = 22.1, p < 0.001. All panels: shading or error bars, 68% confidence interval across animals (N=88, n=1983 sessions).

Across blocks, we found that mice were more liberal (licked more, irrespective of correctness, quantified as a negative bias) in each of the high, compared to neighboring low-reward blocks within an experimental session (Fig. 2A-C). This pattern represents a performance optimization, because bias in high utility was closer to optimal (zero) bias, but not at attention optimization, because bias is not related to sensory signal detection (see **Fig. S2A**). However, sensitivity was also higher in the high-reward compared to low-reward blocks, especially early in each session (**Fig. 2D**-**F**), indicating that feature-based attentional intensity was being adapted to the current task utility. Mice became more conservative and less sensitive across the session duration (**Fig. 2A,C,D,F**), probably resulting from effects of fatigue and/or satiety, both of which would progressively decrease the relative utility of performing the task as the session progresses. Mirroring the patterns in bias and sensitivity, reaction times were substantially shorter in high-reward blocks than in low-reward blocks, and gradually increased as the experimental session progressed (Fig. 2G**-I**). In sum, increases in task utility boosted both task engagement, as indicated by the shift towards a liberal bias, as well as feature-based attentional intensity, as indicated by increased sensitivity and decreased RT in high utility.

To understand how quickly mice could deploy attentional intensity, we next sought to determine the within-block time course of these block-based behavioral changes. Because the reward context was not cued, the first reward in a block provided a strong and unambiguous indication that a transition in utility had occurred. Because the only cue the animals had about which reward block they were in was the observed reward size, we analyzed time-dependencies within each block, aligned to the first hit trial after each block transition (**Fig. 2A**,**D,G,J**). After the first large reward in the high-reward block, bias immediately switched from conservative to liberal, compared to just before the switch, and did not subsequently change during the block (Fig. 2B). In contrast, when switching from high to low reward, bias immediately went from neutral to conservative and then became even more conservative by a further 151.7% with a time constant of 15 trials (see Methods for details). These results show that mice could increase their engagement, rapidly, in high utility, but spent many trials decreasing their engagement in low utility.

After the first high reward in the high-reward block, sensitivity immediately increased by 28.3% compared to just before the switch, and then increased by a further 2.5% with a time constant of 5 trials (Fig. 2E). When switching from high to low rewards, sensitivity immediately increased by 35.2% (likely due to residual high attention from the previous high-reward block and then decreased by 37.9% with a time constant of 15 trials, mirroring the trends for bias. Thus, we observed a hysteresis effect, similar to what has previously been observed in monkeys (Ghosh & Maunsell, 2021). Specifically, mice updated their behavior faster when switching from low to high reward, than the other way around, particularly for sensitivity. This hysteresis indicates a heightened urgency when task utility is detected to have increased.

An animal or human optimally performing a task should allocate cognitive resources in a way that maximizes the rate of returns. We thus wondered if mice collected more rewards in the context of high task utility. This is not trivially so in the sustained-attention value task; unlike in standard go/no-go tasks, in our quasi-continuous task a noise stimulus always precedes each signal stimulus. Thus, a liberal bias, leading to more false alarms, lowers the probability that signals are presented and thus lowers reward rate (**Fig. S2A**). Mice did ignore the noise and detected the signal on a larger fraction of trials in the high-reward compared to low-reward blocks and thus collected more rewards (**Fig. 2J**-**L**). Mice on average sustained a stable reward probability of 0.278 ± 0.006 across the entire session during the high-reward blocks (odd numbered), while reward probability declined monotonically from 0.256 ± 0.008 in the first low-reward block (block 2) to 0.210 ± 0.009 in the last (block 6; **Fig. 2K**,**L**). Reward probability exhibited similar hysteresis effects as bias and sensitivity (Fig. 2K). Thus, mice updated all aspects of their behavior more quickly when switching from low to high reward than when switching in the other direction. We verified that our results were robust to specifics regarding trial selection (**Fig. S2H-K**; Methods) and were not confounded by effects of time-on-task (**Fig. S2L-O**; Methods). We ruled out the possibility that animals adjusted their behavior to the previous outcome (reward versus non-reward) rather than to block-wise changes in task utility. After a previous reward, mice were indeed more liberal, more sensitive, and faster, but task utility had similar effects as before even when accounting for previous trial outcome (**Fig. S2P-S**).

In a parallel analysis, we split each trial into its noise and signal epochs (only noise for false alarms) and then performed a multiple logistic regression of response (lick versus no lick) on signal (noise versus signal), time-on-task (trial number), previous outcome (reward versus no reward) and task utility (low versus high) (Fig. 3A; Methods). While doing so, we observed the same behavioral patterns as in the above stratification analyses. Namely, we observed a negative effect of time-on-task and positive effects of previous outcome and task utility on overall responsiveness (closely related to bias; Fig. 3B) and a negative effect of time-on-task and positive effects of previous outcome and task utility on signal-selective responsiveness (closely related to sensitivity; Fig. 3C).

**Figure 3.**
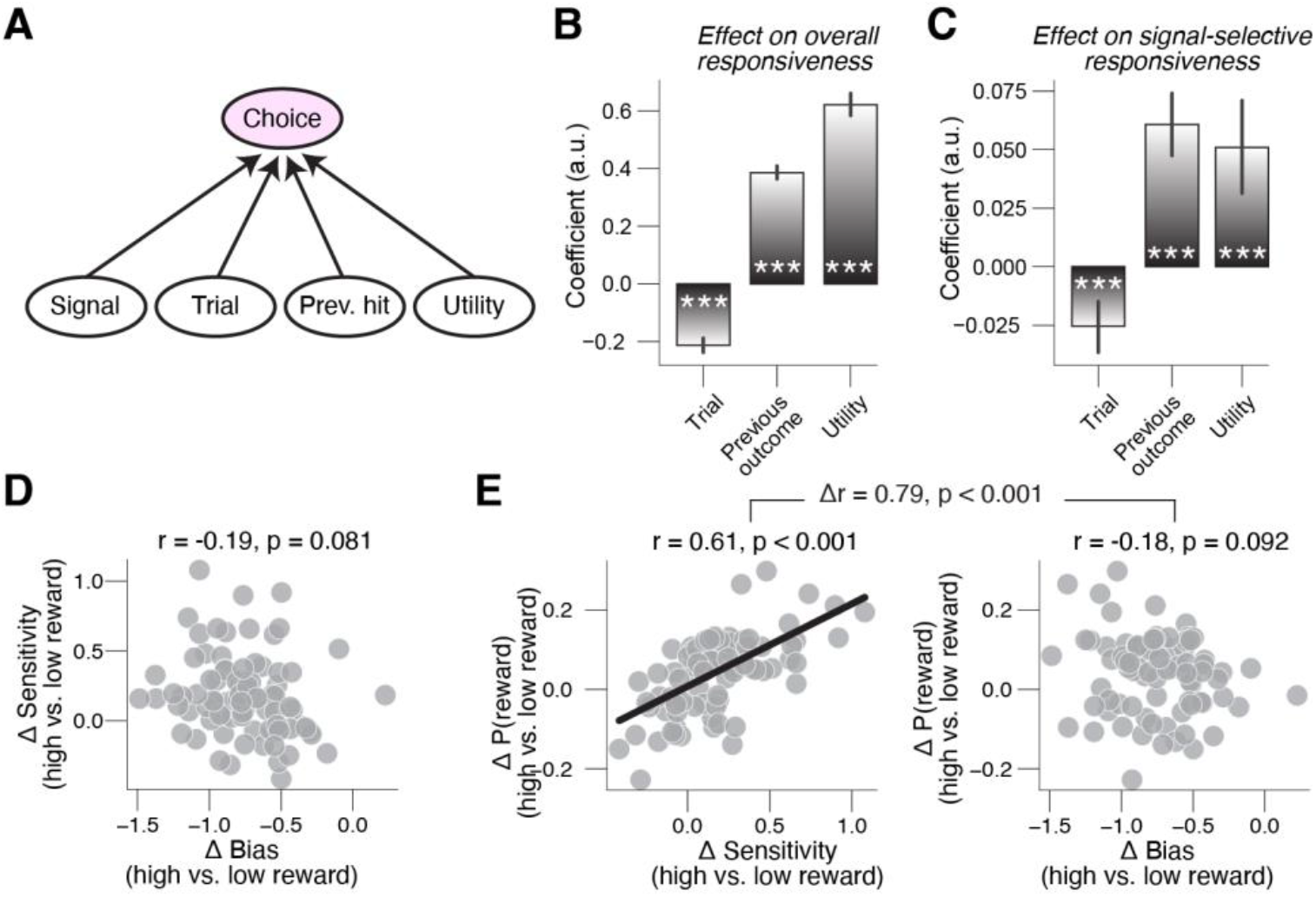
Adaptive allocation of attentional intensity is not an artifact of previous trial outcome and is apparent in patterns of individual difference. **(A)** Schematic of logistic multiple regression of choice on signal [present / absent], trial number, previous outcome [reward / no reward], and utility [low / high] (Methods). **(B)** Fitted coefficients from multiple logistic regression model (panel I; Methods), capturing the effects of time-on-task, previous hit, and utility on overall responsiveness (closely related to bias). Stats, Wilcoxon signed-rank test; ***, p < 0.001; error bars, 68% confidence interval across animals (N=88, n=1983 sessions). **(C)** As B, but for interaction effects between each factor and signal, capturing the effects of time-on-task, previous hit, and utility on signal-selective responsiveness (closely related to sensitivity). **(D)** Change in sensitivity (high vs low task utility) plotted against change in bias. Every data point is a unique session. **(E)** Left: Change in reward probability (high vs low task utility) plotted against change in sensitivity (high vs low task utility). Every data point is a unique session. Right: As left, but for change in bias on the x-axis.

Finally, we asked if the block-wise changes in reward probability were predicted by behavioral adjustments in bias or sensitivity. An individual difference analysis favored the latter. In the first place, the overall (**Fig. S3A**) and block-based (Fig. 3D) changes in bias and sensitivity were not positively correlated at the per-mouse level, suggesting separate underlying processes. Critically, utility-driven shifts in reward probability were strongly predicted by matched shifts in sensitivity, but not bias (Fig. 3E). We conclude that mice increase their feature-based attentional intensity during periods of high task utility; when utility is high, they are more sensitive in discriminating signals from noise, are faster at doing so, resulting in the collection of more rewards.

### Mid-level pupil-linked arousal is optimal for fast and accurate attention-task performance

We previously showed that optimal signal-detection behavior in a simple tone-in-noise detection task occurred at intermediate levels of spontaneously fluctuating pupil-linked arousal (McGinley, David, et al., 2015). Based on these results, we sought to test the hypothesis that the elevated task engagement (liberal bias shift) and attentional intensity (increased d’) we observed during high task utility were partly mediated by stabilization of arousal near its optimal state. To do so, we first performance in the task depended on arousal.

We quantified arousal as the diameter of the pupil (see Fig. 1B) measured immediately before each trial. On trials characterized by a mid-size pre-trial pupil size, we observed the smallest bias, highest sensitivity, shortest RTs and highest reward probability (**Fig. 4A**-**D**; Methods). We defined the optimal level of arousal as the pre-trial baseline pupil size bin with maximal reward probability (green vertical line in Fig. 4D), which also was the arousal-defined bin for which bias was minimal, sensitivity was maximal, and RTs were short (dashed green lines in **Fig. 4A**,**B**,**C**). Across animals, this optimal pre-trial baseline pupil size was on average 27.5% of its maximum. Hit rates and false alarm rates also peaked at this pupil size (**Fig. S4D-E**), and similar patterns were observed early and late in blocks and across learning (**Fig. S4L-M**).

**Figure 4.**
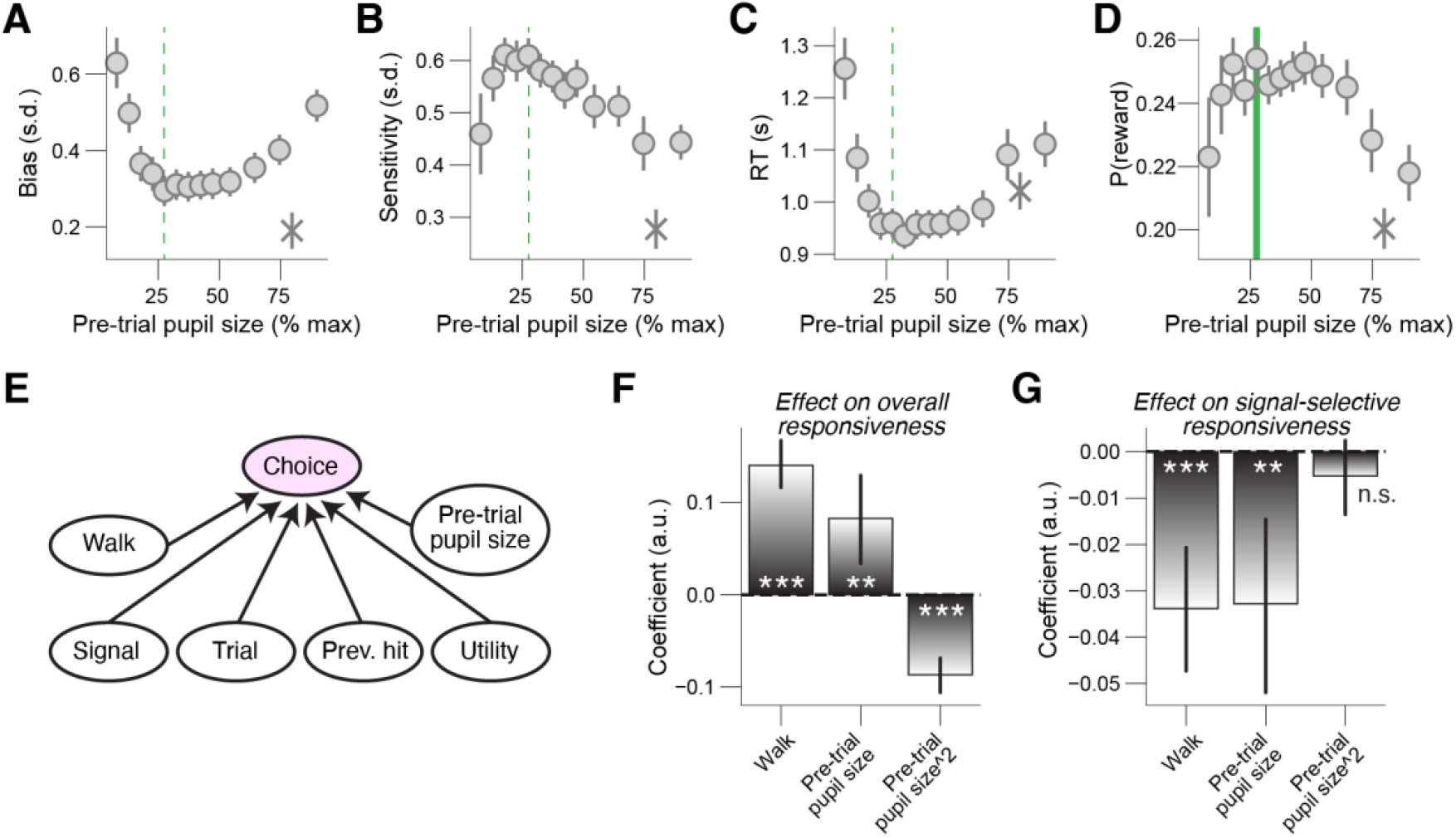
Optimal performance occurs at intermediate levels of pupil-linked arousal. **(A)** Relationship between pre-trial pupil size and bias (irrespective of task utility; Methods). A 1^st^ order (linear) fit was superior to a constant fit (F_1,12_ = 12.7, p = 0.004) and a 2^nd^ order (quadratic) fit was superior to the 1^st^ order fit (F_1,12_ = 7.8, p = 0.016; sequential polynomial regression; Methods). Asterisk, walking trials (Methods). Dashed green line, optimal arousal state (maximum probability peaks; see panel D). **(B)** As A, but for sensitivity. 1^st^ order fit: F_1,12_ = 16.6, p = 0.002; 2^nd^ order fit: F_1,12_ = 0.7, p = 0.430. **(C)** As A, but for RT. 1^st^ order fit: F_1,12_ = 3.2, p = 0.101; 2^nd^ order fit: F_1,12_ = 5.7, p = 0.034. **(D)** As A, but for reward probability. 1^st^ order fit: F_1,12_ ∼ 0.0, p = 0.926; 2^nd^ order fit: F_1,12_ = 7.2, p = 0.020. Green line, optimal arousal state (maximum probability peaks). **(E)** Schematic of logistic multiple regression of choice on signal [present / absent], trial number, previous hit [hit / no hit], utility [low / high], pre-trial walking [still / walk] and pre-trial pupil size (Methods). **(F)** Fitted coefficients from multiple logistic regression model (panel E; Methods), capturing the effects of pre-trial walking and pre-trial pupil size on overall responsiveness (closely related to bias). Stats, Wilcoxon signed-rank test; **, p < 0.01; ***, p < 0.001. **(G)** As F, but for interaction effects between each factor and signal, capturing the effects of pre-trial walking and pre-trial pupil size on signal-selective responsiveness (closely related to sensitivity). All panels: error bars, 68% confidence interval across animals (N=88, n=1983 sessions).

Locomotor status is another widely used marker of behavioral state (McGinley, David, et al., 2015; Polack et al., 2013). Mice were considerably more liberal and less sensitive on trials associated with pre-trial walking (asterisks in **Fig. 4A**,**B**) and reward probability was at its lowest (Fig. 4D). These results suggest that responses during walking were quite random (not signal-related), and thus that attentional intensity was very low during locomotion. We extended our logistic regression model by including three additional predictors: walking (walk versus still), pupil-linked arousal (pre-trial pupil size) and pre-trial pupil size raised to the power of two (Fig. 4E; Methods). Results in the logistic regression recapitulate the stratification-based effects of pupil (**Fig. 4F**,**G** and **Fig. S4F**,**G**). Using the logistic regression analysis approach we also verified that none of our results can be explained by whether mice walked on the previous trial (**Fig. S4H,I**).

In sum, on trials characterized by a mid-size pre-trial pupil size, we observed the smallest bias, highest sensitivity, shortest RTs and highest reward probability. We conclude that mid-level pupil-linked arousal is the optimal state for fast and accurate feature-based sustained attention.

### Utility-related performance improvements are partially mediated by stabilization of pupil-linked arousal near optimality

Having identified the optimal state for performance of the task, we next determined whether mice spent more time in this optimal state during periods of high task utility. The pre-trial pupil size was overall smaller in the high-reward compared to low-reward blocks (**Fig. 5A**-**C** and **Fig. S5F-H**). Likewise, mice walked less in the high-reward compared to low-reward blocks (**Fig. 5D**-**F**). These results suggest an overall average reduction in arousal in high utility context. Since the optimal pupil-linked arousal level of 27.5% was below the average level for all block, which ranged from 37% to 39%, this reduction in arousal was a shift towards optimality. To capture how close the animals’ arousal state on each trial was to the optimal level, we computed the absolute difference between each pre-trial’s pupil size and the optimal size (Methods). The distance from optimality was lower in the high utility context (**Fig. 5G**-**I**). Furthermore, as the session progressed, their arousal state became less optimal; but importantly, mice maintained their arousal closer to the optimal state in the high-reward compared to the low-reward blocks across the session (**Fig. 5G**,**I**).

**Figure 5.**
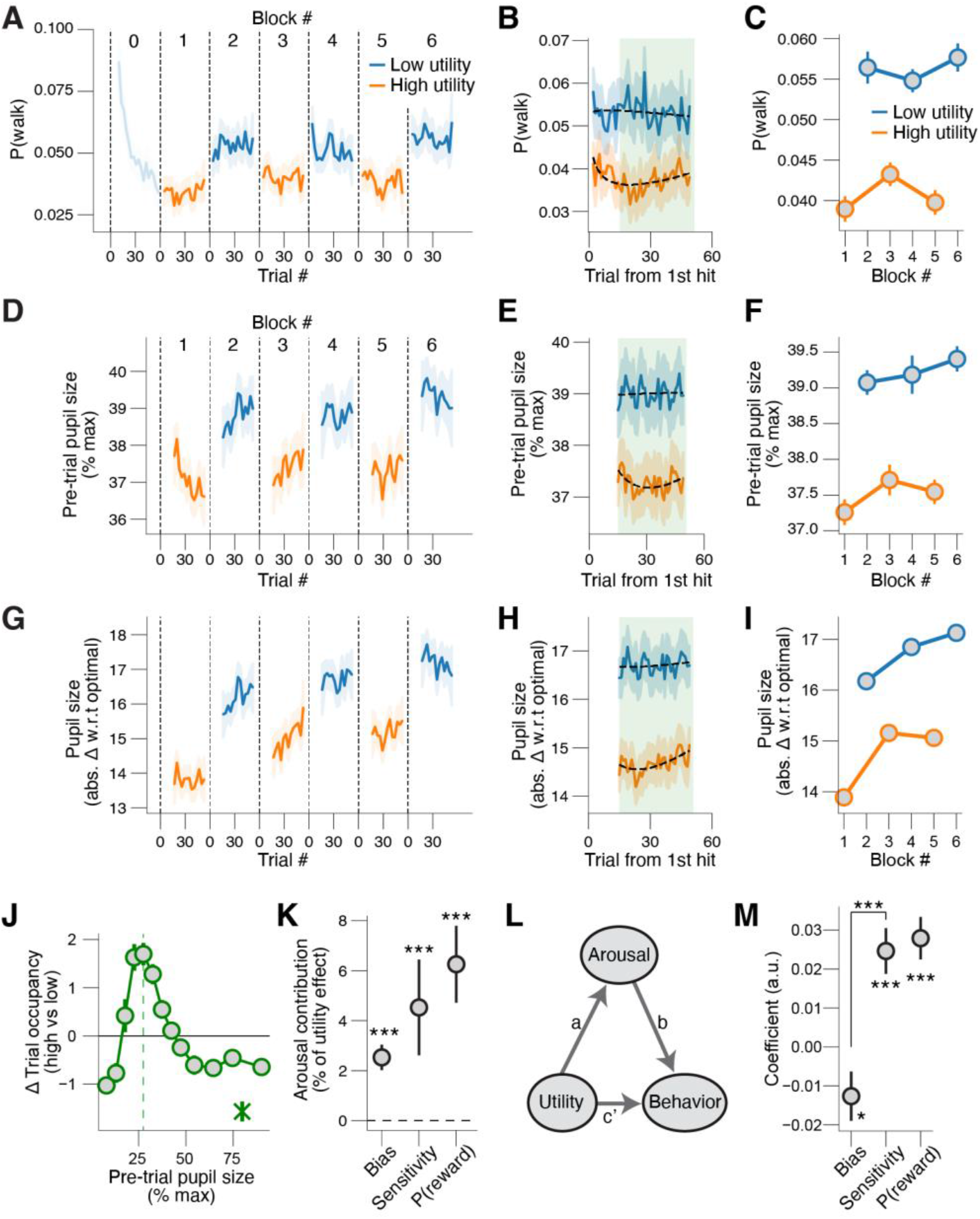
Mice regulate their pupil-linked arousal towards optimality when task utility is high. **(A)** Walk probability (Methods) across low-reward and high-reward blocks in each experimental session, locked to first hit in block. Data from the first block of each session (low utility; termed block ‘0’) was excluded from all analyses, as mice spent this block becoming engaged in the task (see also Fig. 2A,D,G,J; Methods). **(B)** As A, but collapsed across blocks of same reward magnitude. The green shaded area indicates the trials used when pooling data across trials within a block (e.g. panel C). **(C)** As A, but collapsed across trials within a block. Stats, 2-way repeated measures ANOVA (factors task utility [high vs. low] and time-on-task [early, middle, late]); main effect task utility: F_1,87_ = 58.5, p < 0.001; main effect time-on-task: F_2,174_ = 0.3, p = 0.710; interaction effect: F_2,174_ = 3.3, p = 0.039. **(D-F)** As A-C, but for pre-trial pupil size (time-on-task and previous hit regressed out; see Methods; see Fig. S5F-L for results based on raw pre-trial pupil size). Main effect task utility: F_1,87_ = 35.5, p < 0.001; main effect time-on-task: F_2,174_ = 1.7, p = 0.187; interaction effect: F_2,174_ = 1.0, p = 0.376. **(G-I)** As A-C, but for distance of pupil size from the optimal level. Stats, 2-way repeated measures ANOVA (factors task utility [high vs. low] and time-on-task [1, 2, 3]); main effect task utility: F_1,87_ = 125.6, p < 0.001; main effect time-on-task: F_2,174_ = 28.5, p < 0.001; interaction effect: F_2,174_ = 2.7, p = 0.070. **(J)** Change in trial density after increases in task utility, separately for pupil-defined arousal states; asterisk, walking trials. **(K)** Percentage of the purely arousal-predicted shift in behavior compared to the total shift in behavior after changes in task utility (see main text). Stats, Wilcoxon signed-rank test; ***, p < 0.001. **(L)** Schematic of mediation analysis of task utility to behavior, via distance w.r.t. optimal measure (Methods). Arrows, regressions; coefficient *a* × *b* quantifies the ‘indirect’ (mediation) effect; coefficient *c’* quantifies the ‘direct effect’. **(M)** Fitted regression coefficients of the indirect path (*a* × *b*; mediation). Stats, Wilcoxon signed-rank test; *, p < 0.05; ***, p < 0.001. All panels: error bars, 68% confidence interval across animals (N=88, n=1983 sessions).

We sought to determine if the changes in state in high utility context were characterized by uniform down-regulation of arousal, or by stabilization near the optimal level. To test this, we first compared the distributions of state occupancy between low and high reward contexts. Strikingly, mice spent less time in both the low and high arousal states, both of which are suboptimal (see Fig. 4A**-D**) and upregulated a narrow range of states around optimality (Fig. 5J and **Fig. S5A**).

We next tested whether changes in task utility were more associated with changes in the average pre-trial pupil size or with deviations of the pre-trial pupil size from the optimal size. To do so, we first compared the effect sizes of the main effects of task utility on both pupil-linked arousal measures. The partial η^2^ was 0.33 for pre-trial pupil size and 0.55 for its distance from optimal, indicating a larger effect size for distance from optimal compared to raw pre-trial size. Secondly, we performed a logistic regression of block-wise reward magnitude (indicated as 0 or 1) on either z-scored block-wise pre-trial pupil size or its distance from optimal. The fitted coefficients were negative in both cases, but significantly more so for the measure of distance from optimality (**Fig. S5B**). Thus, during heightened task utility, mice do not stereotypically downregulate their arousal state, but instead up-and down-regulate their arousal closer to its optimal level.

Having observed and quantified that epochs of high task utility are associated with both a more optimal pupil-linked arousal state and increased behavioral performance, we wondered to what extent the arousal stabilization contributed the utility-related performance effects. We addressed this with two complementary approaches. In a first approach, we first computed the pupil-linked and walk-related arousal-predicted behavioral performance in the high-reward and low-reward blocks by plugging each trial’s pre-trial pupil size into the previously observed relationship between arousal bins and behavior (see Fig. 4A**-D**; Methods). We then computed the difference in behavioral performance between the high-reward and low-reward blocks that would have occurred solely as a result of the differing pupil-linked and walk-related arousal states between utility contexts. This result showed that a small, but highly significant, portion of the observed utility-based shift in performance could be accounted for solely by the shifts in pupil-linked and walk-related state occupancy (Fig. 5K). We observed similar effects when only considering pupil-linked arousal and not whether mice walked or remained still (**Fig. S5C**).

In a second approach, we tested for statistical mediation (Baron & Kenny, 1986) of arousal in the apparent effect of task utility on the different performance metrics (**Fig. 5L,M**). We found that (the indirect path of) block-wise increases in task utility predicting block-wise decreases in distance from the optimal pupil-linked arousal state, in turn driving block-wise increases in sensitivity and reward probability, partially mediate the apparent effect of task utility on behavior (Fig. 5M). We observed similar effects when only considering trials during which mice did not walk (**Fig. S5D**) and when using walk probability instead of distance from the optimal arousal state as a mediator (**Fig. S5E**). Finally, we observed similar effects when repeating all analyses without having regressed out effects of time-on-task and previous outcome from the pre-trial pupil size measures (**Fig. S5F-L**; Methods). Taken together, we conclude that regulating pupil-linked arousal towards an optimal level partially implements the adaptive behavioral adjustments that match attentional intensity to its utility.

### Accumulation-to-bound modeling of decision-making in the sustained attention-value task

Because the signal stimulus in our task was a high-order spectro-temporal statistic that emerged at unpredictable times in ongoing noise, correct detection required accumulation across time of partial evidence. Consistent with this perspective, reaction times were overall long and variable (see **Fig. 6C,D**). We therefore applied accumulation-to-bound sequential sampling modeling, as is commonly used with similar tasks in the primate visual system (Gold & Shadlen, 2007), or for auditory click discrimination in rodents (Brunton et al., 2013). It has been shown that rodents can perform acoustic evidence accumulation (Brunton et al., 2013; Hanks et al., 2015), but it is not known if and how evidence accumulation is shaped by task utility or arousal.

**Figure 6.**
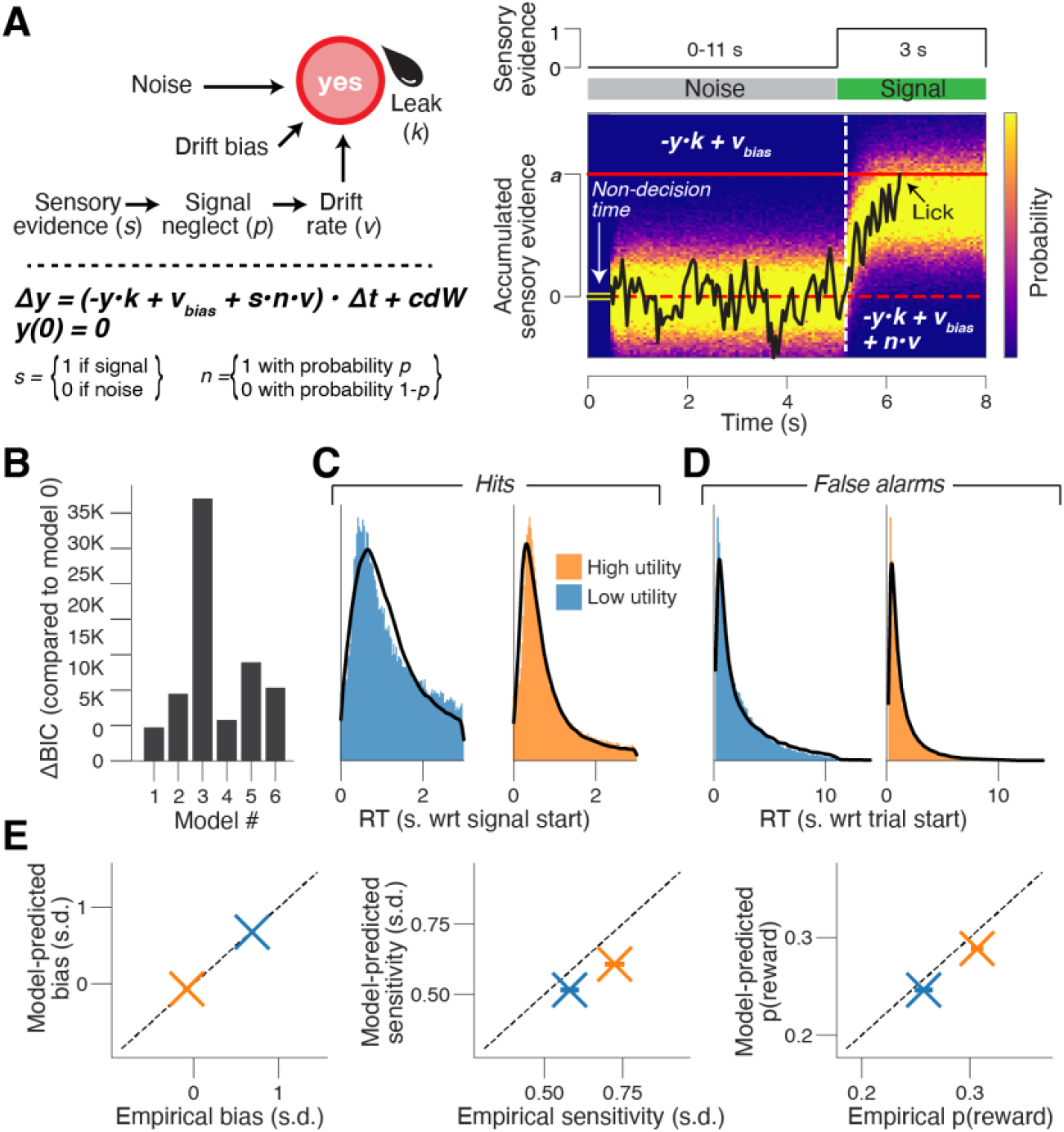
A leaky accumulation-to-bound model accounts for behavior in the sustained-attention value task. **(A)** Left: Schematic of bounded accumulation model accounting for the fraction of go-responses and their associated reaction times (RT). Right: The decision-dynamics were governed by leak (*k*), drift bias (*v_bias_*) and gaussian noise during the tone cloud and additionally signal neglect probability (*n*) and drift rate (*v*) during the signal stimuli. The decision terminated at bound height *a.* **(B)** We compared the Bayesian Information Criterion (BIC) between seven models. The BIC for the winning model was used as a baseline. Lower BIC values indicate a model that is better able to explain the data, taking into account the model complexity; a ΔBIC of 10 is generally taken as a threshold for considering one model a sufficiently better fit. **(C)** RT distribution for correct responses (hits) in the low-reward (left) and high-reward (right) blocks. Black line, model fit. **(D)** As C, but for incorrect responses (false alarms). **(E)** Model-predicted bias (left), sensitivity (middle) and reward probability (right) in the low-reward and high-reward blocks plotted against the empirical estimates. Dashed line, identity line. Panels C-E: pooled data across animals (N=88) and sessions (n=1983).

Widely used accumulation-to-bound models describes the complete accumulation (i.e., without forgetting) of noisy sensory evidence as a decision variable that drifts to one of two decision bounds. Crossing a decision bound triggers a response, specifying a reaction time (Bogacz et al., 2006; Brody & Hanks, 2016; Laming, 1968; Ratcliff & McKoon, 2008). When the evidence is stationary (i.e. its summary statistics are constant across time), as occurs in typical go/no-go or two-alternative forced choice tasks, this model produces the fastest decisions for a fixed error rate (Bogacz et al., 2006). In our task, however, like perceptual decisions in most natural settings, the relevant evidence is not stationary. In this case, complete integration is suboptimal, because it results in an excessive number of false alarms due to integration of pre-signal noise (Ossmy et al., 2013). We thus used a computational model of the decision process based on leaky (i.e. forgetful) integration to a single decision bound (Usher & McClelland, 2001); see Methods for details).

Our winning model contained six main parameters (Fig. 6A**-B**; **Fig. S6A-L**; Methods), the choice of which was motivated by the design of the task and by general patterns observed in the behavior. The six parameters were: (i) bound height, which determines how much evidence needs to be accumulated before committing to a go-response; (ii) non-decision time, which captures the combined duration of pre-decisional evidence encoding and post-decisional translation of choice into motor response; (iii) leak, which controls the timescale of evidence accumulation; (iv) drift bias, which is an evidence independent constant that is added to the drift; (v) mean drift rate, which controls the efficiency of accumulation of the relevant sensory feature (temporal coherence in this case); and (vi) signal neglect probability, which is the fraction of signal epochs for which the task-relevant sensory evidence is not accumulated. The fitted model accounted well for the behavior in the sustained-attention value task, making accurate predictions for both RTs (Fig. 6C**-D**), as well as utility context-based bias, sensitivity, and reward rate (Fig. 6E).

We considered six plausible alternative models (see Methods), including one with variable bound height, which provided worse fits, both qualitatively (Fig. 6B) as well as quantitatively (**Fig. S6**). Two alternative models established the necessity including the signal neglect probability parameter: this was necessary to account for the observation that correct responses (hits) were faster than expected for their frequency of occurrence (**Fig. S6E-H**). Likewise, a drift bias was necessary to account for fast errors (false alarms), especially in the low reward blocks (**Fig. S6I-L**).

### Attention-based decision computations improve during high task utility

We next sought to dissociate distinct elements of the decision-making process underlying the observed effects of task utility on our SDT metrics of behavior (see Fig. 2). Because there were prominent effects of time-on-task, previous trial outcome and block-based task utility in our overt behavioral measures (see Fig. 2 and Fig. 3), we first fitted the full model separately per block number and previous outcome; only bound height was fixed across conditions (see Methods). We found that the drift bias and drift rate were higher, and the leak and signal neglect probability were lower, in the high-reward vs. low-reward blocks (Fig. 7B**-E**). There was no significant effect of task utility on non-decision time (Fig. 7A). The leak and signal neglect probability increased and drift bias decreased with time-on-task (**Fig. 7B,C**,**E**). These effects were similar when additionally accounting for previous outcome (**Fig. S7A-E**). The main pattern of effects did not depend on the specifics of the model: for each of the alternative models, leak and signal neglect probability were lower in the high-reward compared to low-reward blocks, and drift bias and drift rate were higher (**Fig. S7U-Z**).

**Figure 7.**
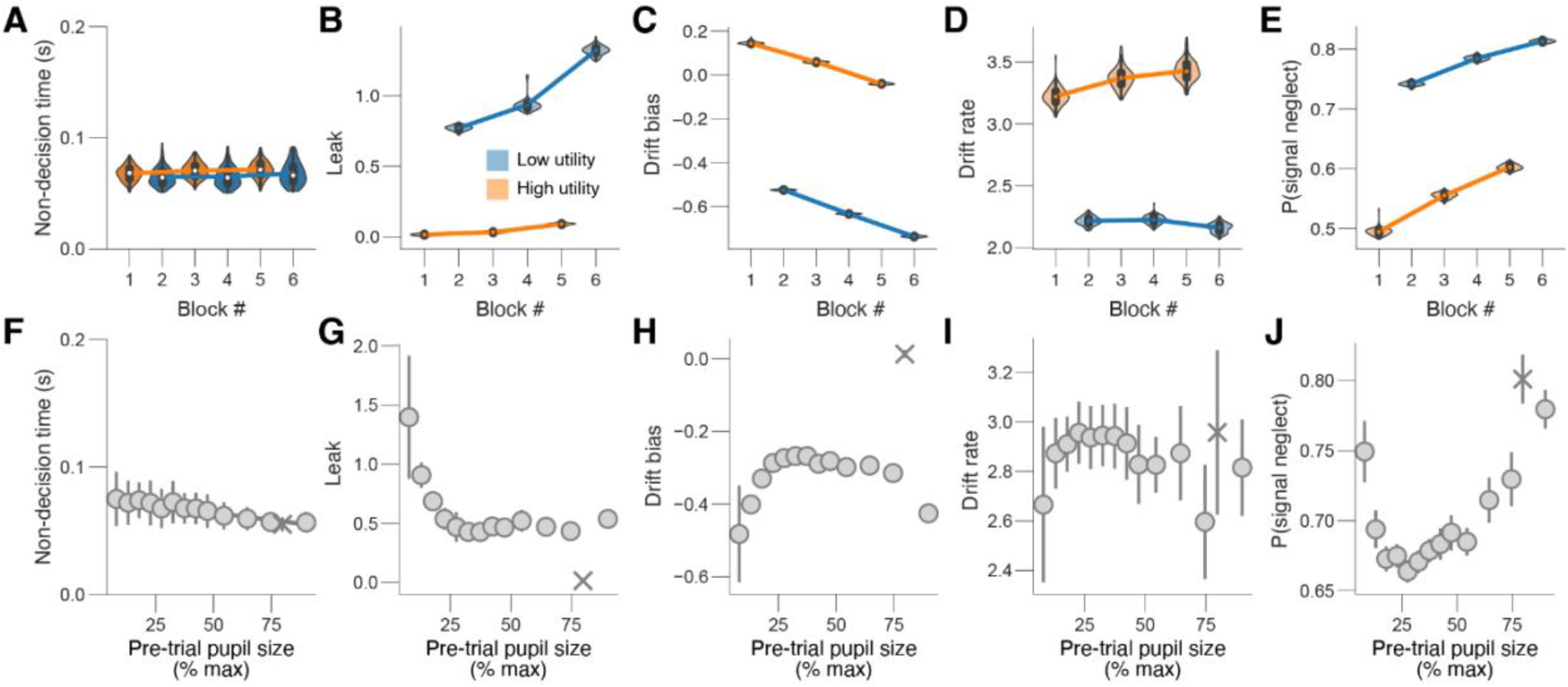
Task utility and pupil-linked arousal impact intersecting aspects of the decision computation. **(A)** Fitted non-decision time estimates (kernel density estimate of 100 bootstrapped replicates) separately per block number. Main effect task utility (fraction of bootstrapped parameter estimates in the low-reward blocks higher than in the high-reward blocks): p = 0.32. Main effect time-on-task (fraction of bootstrapped parameter estimates in the first two blocks higher than in the last two blocks): p = 0.41. **(B)** As A, but for leak. Main effect task utility: p < 0.01. Main effect time-on-task: p < 0.01 **(C)** As A, but for drift bias. Main effect task utility: p < 0.01. Main effect time-on-task: p < 0.01. **(D)** As A, but for drift rate. Main effect task utility: p < 0.01. Main effect time-on-task: p = 0.46. **(E)** As A, but for signal neglect probability. Main effect task utility: p < 0.01. Main effect time-on-task: p < 0.01. **(F)** Fitted non-decision estimates (100 bootstrapped replicates) separately per arousal state (same pupil size defined bins as in Fig. 4A-D; irrespective of task utility; Methods). A 1^st^ order (linear) fit was superior to a constant fit (F_1,12_ = 128.8, p < 0.001) and a 2^nd^ order (quadratic) fit was not superior to the 1^st^ order fit (F_1,12_ = 2.2, p = 0.168; sequential polynomial regression; Methods). Asterisk, walking trials (Methods). **(G)** As F, but for leak. 1^st^ order fit: F_1,12_ = 8.9, p = 0.012; 2^nd^ order fit: F_1,12_ = 5.4, p = 0.039. **(H)** As F, but for drift bias. 1^st^ order fit: F_1,12_ = 8.8, p = 0.391; 2^nd^ order fit: F_1,12_ = 7.7, p = 0.017. **(I)** As F, but for drift rate. 1^st^ order fit: F_1,12_ = 0.8, p = 0.378; 2^nd^ order fit: F_1,12_ = 3.1, p = 0.101. **(J)** As F, but for signal neglect probability. 1^st^ order fit: F_1,12_ = 2.2, p = 0.165; 2^nd^ order fit: F_1,12_ = 7.3, p = 0.019.

In sum, increases in task utility resulted in a longer accumulation time constant (lower leak), a higher urge to respond (higher drift bias), more efficient evidence accumulation (higher drift rate) and more reliable evidence accumulation (lower signal neglect probability).

### Mid-level pupil-linked arousal exhibits low signal neglect probability, leak, and drift bias

We next sought to understand how decision computations, as reflected in the accumulation-to-bound modeling, were affect by pupil-linked arousal state. We thus fitted the full model separately for each of the pupil-linked arousal bins. Non-decision time and leak decreased monotonically with arousal (**Fig. 7F,G**; **Fig. S7F,G**), suggesting faster sensory/motor processing and longer-lasting evidence accumulation in higher arousal states. Drift bias, drift rate and signal neglect probability all exhibited U-or inverted-U shaped dependencies on arousal (Fig. 7H**-J**; **Fig. S7H-J**) indicating that the mice more reliably accumulated evidence from, and acted upon, the signal at moderate arousal levels. As with the utility-related effects, we observed similar effects of pupil-linked arousal when repeating the analyses without having regressed out effects of time-on-task and previous outcome from the pre-trial pupil size measures (**Fig. S7K-T**; Methods).

Thus, attention to the relevant feature is maximal at intermediate levels of pupil-linked arousal, and worst during walking.

## Discussion

To efficiently meet their survival needs, organisms must regulate both of what Daniel Kahneman termed the *selective* and *intensive* (or effortful) aspects of attention (Kahneman, 1973). Characterizing the selective aspect of attention has been a cornerstone of systems neuroscience (Carrasco, 2011; Fritz et al., 2007; Maunsell & Treue, 2006), but attentional intensity has received comparatively little scrutiny. Recent work emphasizes the importance of motivational factors in driving attentional intensity (also called ’attentional effort’; Brehm & Self, 1989; Ghosh & Maunsell, 2021; Kurzban et al., 2013; Richter et al., 2016; Sarter et al., 2006; Shenhav et al., 2013). This emphasis is in line with the common experience of ‘paying more attention’ when motivated to do so, for example in a classroom setting when the instructor indicates that forthcoming material will be on a final exam. In line with these ideas, heightened reward expectation increases perceptual sensitivity and reduces reaction times in humans and non-human primates (Engelmann & Pessoa, 2014; Ghosh & Maunsell, 2021; Locke & Braver, 2008). Parallel work has focused on the effects of arousal level on task performance, typified by the Yerkes-Dodson inverted-U dependence of performance on arousal level (Yerkes & Dodson, 1908; McGinley, David, et al., 2015).

We here developed a feature-based sustained attention task for head-fixed mice with nonstationary task utility and assessed the role of arousal fluctuations in regulating attentional intensity. By applying signal detection theoretic and accumulation-to-bound modeling to a large behavioral and physiological dataset we showed that in contexts of high task utility, mice: (i) collect more rewards, (ii) accumulate perceptual evidence more efficiently, reliably, and across longer timescales, iii) suppress task-irrelevant locomotor behavior, and (iv) stabilize their pupil-linked arousal state closer to an optimal level.

### Nonstationarity in task performance and its causes

Growing evidence supports that neural computation and behavioral performance are not stationary within a session, due to ongoing fluctuations in internal state. State-dependence of neural activity has been observed in primary sensory cortices (Goris et al., 2014; McGinley, David, et al., 2015; Musall et al., 2019; Nestvogel & McCormick, 2022) and sensory-guided behavior (McGinley, David, et al., 2015). More recently, spontaneous shifts between engaged, biased, and/or disengaged states have been inferred from behavior (Ashwood et al., 2022; Hulsey et al., 2023; Weilnhammer et al., 2023). However, the underlying causes and behavioral functions of these non-stationarities are not unclear.

One known major source of non-stationarity in behavior is time-on-task, wherein fatigue and/or satiety result in declining task utility, and associated performance, across a behavioral session. In our results, mice sustained their highest level of performance – encapsulated in the reward probability – across the three high-reward blocks interspersed across a long-lasting and difficult sustained-attention task. Performance was comparably high in only the first low-reward block and then declined precipitously in subsequent low-reward blocks. Our accumulation to bound modeling added that it is mainly the leak, drift bias, and probability of signal neglect that are responsible for the large perceptual performance difference across time on task. A parsimonious interpretation of these findings is that early in the session mice were hungriest and least fatigued, and thus highly motivated to work for any reward. Later in the session, when more satiated and fatigued, only large rewards were sufficient to motivate them to increase their attentional intensity (Hernández-Navarro et al., 2021). An additional factor, which may have increased performance in the low reward and blunted an even larger effect of attentional effort allocation, is that in our task mice needed to keep performing in the low reward context at a sufficiently high level to detect the switch from low to high-reward block. Thus, their capacity to temporally allocate attention is probably even higher than it appeared in our results.

### Motivated shifts in attentional intensity/effort

Task utility-based shifts in behavior, but not attention, have been observed in rodents (Reinagel, 2021; A. Y. Wang et al., 2013). Other than in humans, the regulation of attentional intensity by utility has only been studied in monkeys (Ghosh & Maunsell, 2021). Rodents can match their rate of learning to the statistics of a dynamic environment (Grossman et al., 2022), perform cost-benefit analysis (Reinagel, 2021) and adapt their response vigor to task utility (A. Y. Wang et al., 2013). A recent study demonstrated that mice can consider their own information processing limitations to adaptively allocate the selectivity of their visual attention (Grujic et al., 2022), although changes in overall task utility were not studied.

Our results show that, in the presence of nonstationary overall task utility, but stable selectivity requirements, both task engagement (as indicated by choice bias) and attentional intensity (as indicated by increased d’) increased when task utility was high. In our accumulation-to-bound modeling, the dependencies of leak, drift rate, and signal neglect probability on task utility all support the conclusion that attentional intensity increased in the high-reward blocks. The higher drift rate and lower signal neglect probability in the high, compared to low, reward blocks contributed to a positive effect of task utility on sensitivity and reward probability. The lower leak in high reward context indicates more sensory stimulus engagement, in the form of integrating the stimulus for a longer time, which is another logical component of attentional intensity. Low leak may, or may not, be optimal for reward harvesting, however, because more noise is accumulated with a lower leak. Too low of a leak would be an example of paying too much attention. Future study could use the sustained attention value task to explore this interesting distinction between optimal leak versus excess sensory evidence accumulation.

### The Yerkes-Dodson inverted-U model and motivated attention

Our results add to the growing evidence for an inverted-U, three-state model for the role of pupil-linked arousal in behavioral performance (Beerendonk et al., 2023; Hulsey et al., 2023; McGinley, David, et al., 2015; McGinley, Vinck, et al., 2015; Schriver et al., 2018; Yerkes & Dodson, 1908). As in our previous work using a perceptually simpler detection task and stationary task utility (McGinley, David, et al., 2015), we here found that the optimal state for task performance was not at either extreme of arousal, but rather was at mid pupil-linked arousal level.

Small pupil-size has been associated with sharp-waves in the hippocampus and slow oscillations in neocortex (McGinley, David, et al., 2015), which are classic signatures of low arousal. In our task, compared to mid pupil size, small pre-trial pupil was associated with: conservative SDT bias, low SDT sensitivity, long RTs, lower reward probability, high leak, conservative drift bias, and high probability of signal neglect; all of these are consistent with drowsiness or some other resting form of disengagement. Large pupil size in our task was associated with conservative SDT bias, low SDT sensitivity, longer RTs, decreased reward probability, increased walking, and a large increase in signal neglect probabiliy; all of these are consistent with a hyper-aroused form of task disengagement. Importantly, mice spent less time in both the under- and over-aroused pupil states when in the high task utility context. Furthermore, arousal-stabilization partially mediates the effect of reward.

### Conceptual models of arousal in attention

Our findings that both pupil-linked and extent of walking were higher in the low-reward blocks indicate that mice did not use the low-reward blocks simply to rest, but instead to engage in alternative, aroused and perhaps exploratory, behaviors. This is in line with a recent observation that lapses in perceptual decisions reflect exploration (Pisupati et al., 2021), which is consistent with the broad notion of an exploration-exploitation tradeoff (Aston-Jones & Cohen, 2005; Gilzenrat et al., 2010; Jepma & Nieuwenhuis, 2011) contributing to the right side of the inverted-U. Our pupil results on the high-arousal right side of the inverted-U are partially consistent with results of a recent study (Grujic et al., 2022), which found that large baseline pupil was associated with task disengagement. However, we observed that large pupil also was associated with a large reduction in accumulation of signal-related sensory evidence (a high signal neglect probability) whereas that study did not find effects of pupil on sensory encoding precision. This difference may result from their use of simple (visual grating) signal detection versus our use of a higher-order acoustic feature (temporal coherence) tracking across time.

Our observed role for pupil-linked arousal in mediating adaptive adjustments in attentional intensity, based on task *utility*, is in contrast with the large literature on pupil dilation as a readout of attentional capacity (also called effort) driven by fluctuations in task *difficulty* (Alnæs et al., 2014; Hess & Polt, 1964; Kahneman et al., 1967; Kahneman & Beatty, 1966; Laeng et al., 2012), including the extensive work on pupil-linked listening effort (Peelle, 2018; Pichora-Fuller et al., 2016). In this literature, the magnitude of the task-evoked pupil response is measured during the stimulus and compared between conditions that differ in difficulty. For example, studies employ multiple levels of speech degradation or memory load (Alnæs et al., 2014; Zekveld et al., 2014). Therein, motivational factors are customarily neglected. This neglect has been widely acknowledged (Pichora-Fuller et al., 2016), but not addressed. In our sustained-attention value task, perceptual difficulty and selectivity (the temporal coherence) were held constant across the session, whereas motivation (driven by task utility) was changed in blocks. Furthermore, we focus on the pre-stimulus, so-called ‘tonic’, pupil-linked arousal measured before each trial, rather than peri-stimulus, so-called ‘phasic’, stimulus-driven pupil dilation (de Gee et al., 2020). Future work is needed to determine the interaction of these complementary arousal functions in behavior, perhaps by combining non-stationarities in both task utility and perceptual difficulty or selectivity.

The specific pattern we observed in the dependence of performance on pupil-linked arousal, and its relation to task utility, likely illustrates both general principles as well as task- and species-specific patterns. For example, in contrast to our findings, a previous human study reported higher pre-trial pupil size during high-reward blocks (Massar et al., 2016). This task was perceptually easy, while our sustained-attention value task was perceptually hard. An extensive literature shows that the relationship between tonic arousal and behavioral performance depends on task difficulty, with higher arousal being optimal for easier tasks (Sörensen et al., 2021; Yerkes & Dodson, 1908). The discrepancy between the findings reported by Massar et al. (2016) and ours might also be due to species difference; perhaps humans in laboratory conditions are on average in a lower arousal state than mice and thus typically sit on the opposite side of optimality. However, this is not the case for all individuals; moving closer to the optimal arousal state after increases in task utility involves either increases or decreases in arousal, depending on one’s starting point (de Gee et al., 2020). The adaptive function of low arousal states (such as for online learning and consolidation) (Pfeiffer, 2020; Squire et al., 2015) and of high arousal states (such as for broadly sampling the environment to observe changes and exploring for alternatives) (Aston-Jones & Cohen, 2005), and self-regulation of sampling of these states during behavior, all require further study.

### Neural substrates for regulation of attention by arousal and motivation

Our results raise the question of which neuromodulatory systems contribute to the effects of arousal on attentional intensity. Our finding that tonic pupil-linked arousal is lower in the low-reward blocks is in line with the adaptive gain theory of LC function (Aston-Jones & Cohen, 2005; Gilzenrat et al., 2010; Jepma & Nieuwenhuis, 2011), but see (Bari et al., 2020), implicating norepinephrine, which plays a major role in pupil control (Breton-Provencher & Sur, 2019; de Gee et al., 2017; Joshi et al., 2016; Reimer et al., 2016; Varazzani et al., 2015). On the other hand, our results are not in line with the idea that increased acetylcholine mediates attention (Hasselmo & McGaughy, 2004; Sarter et al., 2006), but see (Robert et al., 2021). The willingness to exert behavioral control is thought to be mediated by tonic mesolimbic dopamine (Hamid et al., 2016; Niv et al., 2007; A. Y. Wang et al., 2013) and/or serotonin (Gutierrez-Castellanos et al., 2022). However, the willingness to work is likely more related to bias, while attentional intensity is more related to sensitivity. Orexin/hypocretin is also involved in reward processing, and serotonin and orexin/hypocretin have both been implicated in pupil control (Cazettes et al., 2021; Grujic et al., 2023). Thus, possibilities abound, and future work is needed to determine the precise roles of neuromodulatory systems in utility-driven and arousal-regulated allocation of attentional intensity.

Our results suggest a cost-benefit analysis being used to adapt pupil-linked arousal and performance level to an evolving motivational state (Botvinick & Braver, 2015; Shenhav et al., 2013). Neural underpinning of such top-down control of attentional intensity are yet to be elucidated. Orbital frontal cortex (OFC) and dorsal anterior cingulate cortex (dACC) are likely candidate regions since both perform value computations related to optimizing behavior (Akam et al., 2021; Botvinick & Braver, 2015; Shenhav et al., 2013; Tremblay & Schultz, 2000). These structures are strongly connected to sensory cortices and to neuromodulatory nuclei, including LC (Arnsten & Goldman-Rakic, 1984; Aston-Jones & Cohen, 2005; de Gee et al., 2017; Joshi & Gold, 2022; Porrino & Goldman-Rakic, 1982). Impaired frontal regulation of arousal is one of the hallmarks of ADHD (Barkley, 1997) and also plays a role in autism (Zhao et al., 2022) and a wide array of psychiatric disorders (de Lecea et al., 2012; Sander et al., 2015). Future work should determine the top-down influences of frontal cortices on arousal systems supporting attentional intensity, as well as dysregulation in mouse models of neurological disorders.

We here discovered that regulating pupil-linked arousal towards an optimal level partially implements behavioral adjustments that adaptively increase attentional intensity when it is useful. These results suggest that at least a part of the large behavioral and neural variability that can be explained by fluctuations in pupil-linked arousal state serve an adaptive function; states conducive to a particular biological need (i.e. attention to a rewarded stimulus) are upregulated at appropriate times (i.e. when the reward is large).

## Methods

### Animals

All surgical and animal handling procedures were carried out in accordance with the ethical guidelines of the National Institutes of Health and were approved by the Institutional Animal Care and Use Committee (IACUC) of Baylor College of Medicine. A total of 114 animals were trained through to at least 5 sessions of the final phase of the task (see *Behavioral task*). We excluded 26 animals from the analysis (**Fig. S1P-R**) who had less than 5 sessions worth of data per animal after excluding sessions with an overall reward probability (see *Analysis and modeling of choice behavior*) of less than 15%. Thus, all remaining analyses are based on 88 mice (74 male, 14 female) aged 7-8 weeks at training onset. Wild-type mice were of C57BL/6 strain (Jackson Labs) (N=51; 1 female). Various heterozygous transgenic mouse lines used in this study were of Ai148 (IMSR Cat# JAX:030328; N=6; 3 females), Ai162 (IMSR Cat# JAX:031562; N=10, 3 females), ChAT-Cre (IMSR Cat# JAX:006410; N=3; all male), or ChAT-Cre crossed with Ai162 (N=18; 7 females). No differences were observed between genotypes or sexes, so results were pooled. This variety in genetic profile was required to target specific neural circuitries with two-photon imaging; the results of the imaging experiments are not reported, here. Mice received ad libitum water. Mice received ad libitum food on weekends but were otherwise placed on food restriction to maintain ∼90% normal body weight. Animals were trained Monday-Friday. Mice were individually housed and kept on a regular light-dark cycle. All experiments were conducted during the light phase.

### Head post implantation

The surgical station and instruments were sterilized prior to each surgical procedure. Isoflurane anesthetic gas (2–3% in oxygen) was used for the entire duration of all surgeries. The temperature of the mouse was maintained between 36.5°C and 37.5°C using a homoeothermic blanket system. After anesthetic induction, the mouse was placed in a stereotax (*Kopf Instruments*). The surgical site was shaved and cleaned with scrubs of betadine and alcohol. A 1.5-2 cm incision was made along the scalp mid-line, the scalp and overlying fascia were retracted from the skull. A sterile head post was then implanted using dental cement.

### Behavioral task

Each ‘trial’ of the behavior consisted of three consecutive intervals (Fig. 1A): (i) the ‘noise’ or tone cloud interval, (ii) the ‘signal’ (temporal coherence) interval, and (iii) the inter-trial-interval (ITI). The duration of the noise interval was randomly drawn before each trial from an exponential distribution with mean of 5 seconds; this was done to ensure a flat hazard function for signal start time. In most sessions (82.8%), randomly drawn noise durations greater than 11 s were set to 11s. In the remainder of sessions (17.2%), these trials were converted to a form of catch trial, consisting of 14 seconds of noise. Results were not affected by whether sessions included catch trials or not (**Fig. S2T-W**), and thus results were pooled for all further analyses. The duration of the signal interval was 3 s. The duration of the ITI was uniformly distributed between 2 and 3 s, plus an additional second after the last lick during the ITI.

The noise stimulus was a ‘tone cloud’, consisting of consecutive chords of 20 ms duration (gated at start and end with 0.5 ms raised cosine). Each chord consisted of 12 pure tones, selected randomly from a list of semitone steps from 1.5-96 kHz. For the signal stimulus, after the semi-random (randomly jittered by 1-2 semitones from tritone-spaced set of tones) first chord, all tones moved coherently upward by one semitone per chord. The ITI-stimulus was pink noise, which is highly perceptually distinct from the tone cloud. Stimuli were presented free field in the front left, upper hemifield at an overall intensity of 55 dB SPL (root-mean square [RMS]) using an intermittently recalibrated Tucker Davis ES-1 electrostatic speakers and custom software system in LabVIEW.

Mice were head-fixed on a wheel and learned to lick for sugar water reward to report detection of the signal stimulus. Correct-go responses (hits) were followed by either 2 or 12 μL of sugar water, depending on block number. Reward magnitude alternated between 2 and 12 μL in blocks of 60 trials, each. Incorrect-go responses (false alarms) terminated the trial and were followed by a 14 s timeout with the same pink noise ITI-stimulus. Correct no-go responses (correct rejecting the full 14 s of noise) in the sessions that contained catch trails were also followed by 2 or 12 μL of sugar water in some sessions (see above).

Training mice to perform the sustained-attention value task involved three separate phases. In phase 1, the signal was louder than the noise sounds (58 and 52 dB, respectively), ‘classical conditioning’ trials (5 automatic rewards during the signal sounds for the first 5 trials) were included, and there were no block-based changes in reward magnitude (reward size was 5 μL after every hit, across the session). Phase 1 was conducted for four experimental sessions (**Fig. S1A**). In phase 2, we introduced the block-based changes in reward magnitude. Once mice obtained a reward probability higher than 0.25, and the fraction of trials resulting in a false alarm was below 0.5 for two out of three sessions in a row, they were moved up to the phase 3. Phase 2 lasted for 2 -85 (median, 9) experimental sessions (**Fig. S1B**). Phase 3 was the final version of the task, with signal and noise stimuli of equal loudness and without any classical conditioning trials.

In a subset of experiments, the signal quality was systematically degraded by reducing the fraction of tones that moved coherently through time-frequency space. This is similar to reducing motion coherence in the classic random-dot motion task (Newsome et al., 1989). In these experiments, signal coherence was randomly drawn beforehand from six different levels: easy (100% coherence; as in the main task), hard (55-85% coherence), and four levels linearly spaced in between. In the main behavior, coherence ranged from 90-100% on each trial. Performance in the main behavior did not change with coherence level, so results were pooled across coherence levels.

After exclusion criteria (**Fig. S1P-R**), a total of 88 mice performed between 5 and 60 sessions (2100–24,960 trials per subject) of the final version of the sustained-attention value task (phase 3), yielding a total of 1983 sessions and 823,019 trials. A total of 10 mice performed the experiment with psychometrically degraded signals; they performed between 5 and 28 sessions (2083–11,607 trials per subject), yielding a total of 142 sessions and 58,826 trials.

### Data acquisition

Custom LabVIEW software was written to execute the experiments, and synchronized all sounds, licks, pupil videography, and wheel motion. Licks were detected using a custom-made infrared beam-break sensor.

*Pupil size.* We continuously recorded images of the right eye with a Basler GigE camera (acA780-75gm), coupled with a fixed focal length lens (55 mm EFL, f/2.8, for 2/3”; Computar) and infrared filter (780 nm long pass; Midopt, BN810-43), positioned approximately 8 inches from the mouse.

An off-axis infrared light source (two infrared LEDs; 850 nm, Digikey; adjustable in intensity and position) was used to yield high-quality images of the surface of the eye and a dark pupil. Images (504 × 500 pixels) were collected at 15 Hz, using a National Instruments PCIe-8233 GigE vision frame grabber and custom LabVIEW code. To achieve a wide dynamic range of pupil fluctuations, an additional near-ultraviolet LED (405-410 nm) was positioned above the animal and provided low intensity illumination that was adjusted such that the animal’s pupil was approximately mid-range in diameter following placement of the animal in the set-up and did not saturate the eye when the animal walked.

*Walking speed.* We continuously measured treadmill motion using a rotary optical encoder (Accu, SL# 2204490) with a resolution of 8,000 counts/revolution.

### Analysis and modeling of choice behavior

All analyses were performed using custom-made Python scripts, unless stated otherwise.

### Trial exclusion criteria

We excluded the first (low reward) block of each session, as mice spent this block (termed block ‘0’; low reward) becoming engaged in the task (see **Fig. 2A,D**,**G,J** and **Fig. S5A**). We found that a small fraction of trials began during a lick bout that had already started during the preceding ITI. These trials were immediately terminated. These rare ‘false start trials’ (2.5±0.2 % s.e.m. of trials across mice), were removed from further analyses. When pooling data across trials within a block, we always excluded the first 14 trials after the first hit in each block (in both high and low-reward blocks; see also *Time course of behavioral adjustments*, below).

### Model-free behavioral metrics

Reward probability was defined as the fraction of trials that ended in a hit (a lick during the signal). Reaction time on hit trials was defined as the time from signal onset until the response (first lick).

### Signal-detection theoretic modeling

We applied signal detection theory (SDT; Green & Swets, 1966) to compute sensitivity (d’, also called discriminability) and choice bias (c, also called criterion) in the quasi-continuous task. Due to the variable duration of the ‘noise’ stimulus epochs, and the yoking of ‘signal’ and ‘noise’ epochs, care was required in calculating the hit rate (HR) and false alarm rate (FAR) to be used for calculating the SDT measures.

On each trial, if a signal epoch occurred, it was of fixed duration. Therefore, a hit rate could be calculated using the conventional definition:

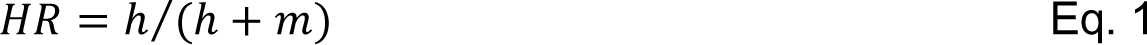

where ‘ℎ’ is the total number of ‘hits’ (signals with a lick response), and ‘𝑚’ is the total number of misses (signals without a lick response).

Because the signal on each trial started at a random time (drawn from an exponential distribution), this HR may depend on the time of the signal start. Therefore, we defined a signal-start time-dependent HR:

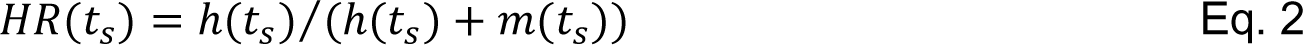

where ‘ℎ(𝑡_𝑠_)’ is the number of ‘hits’ that occurred for signals with a start time of ‘𝑡_𝑠_ ’, and ‘𝑚(𝑡_𝑠_)’ is the number of misses for signals with a start time of ‘𝑡_𝑠_ ’. Because 𝑡_𝑠_ was drawn from a continuous distribution, rather than a discrete list of values, 𝐻𝑅(𝑡_𝑠_) could be empirically estimated in bins of signal start time.

Because licks during the ‘noise’ epoch aborted the trial, resulting in a subsequent signal not being played, the observed signal start-time dependent hit rate corresponds to the following conditional probability:

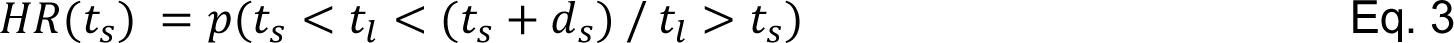

where ‘𝑡_𝑙_ ’ is the time of the first response (lick) and ‘𝑑_𝑠_’ is the duration of the signal. In words, 𝐻𝑅(𝑡_𝑠_) is the probability of licking during the signal, for a signal that starts at time ‘𝑡_𝑠_ ’ given that the mouse did not lick during the preceding noise.

Mirroring the above conditional probability definition of 𝐻𝑅(𝑡_𝑠_), the corresponding conditional probability for time-varying false alarm rate is:

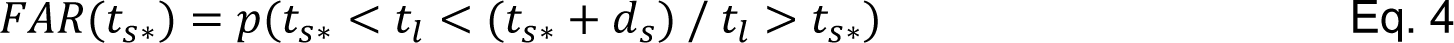

where ‘𝑡_𝑠∗_’ refers to the corresponding signal start time in the ‘𝐻𝑅(𝑡_𝑠_)’ calculation. The ‘*’ is used to indicate that signal had not started yet. In words, ‘𝐹𝐴𝑅(𝑡_𝑠∗_)’ is the time-varying FAR in the appropriate time windows matched to the ‘𝐻𝑅(𝑡_𝑠_)’ function.

Applying Bayes’ rule to 𝐹𝐴𝑅(𝑡_𝑠∗_), and noting that 𝑝(𝑡_𝑙_ > 𝑡_𝑠∗_ / 𝑡_𝑠∗_ < 𝑡_𝑙_ < (𝑡_𝑠∗_ + 𝑑_𝑠_)) = 1, yields:

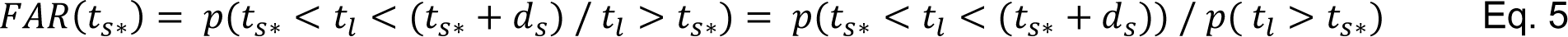

And therefore that:

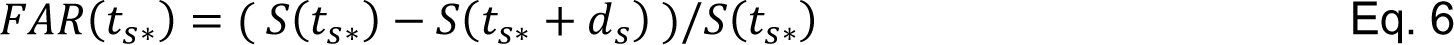

where 𝑆(𝑡) is the survival function for licking during noise. 𝑆(𝑡) could be empirically measured using the Kaplan-Meier estimate, with signal starts treated as right censored false alarm events (Kalbfleisch and Prentice, 2011). Using these time-varying HR and FAR estimates, time-varying SDT measures could be calculated as:

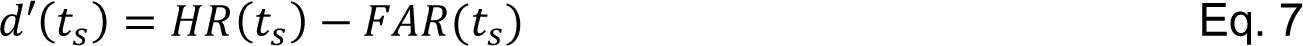

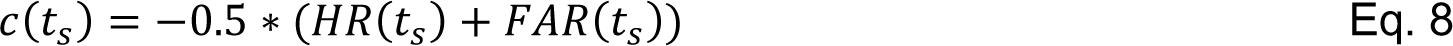

These time-varying SDT estimates can be sampled from the empirical signal start-time distribution and averaged to yield single summary estimates (Macmillan and Kaplan, 1985). All SDT estimates were adjusted by adding 0.5 to each outcome type (hit, miss, correct reject, false alarm) in order to prevent infinite values resulting ‘0’ or ‘1’ for HR or FAR (Hautus, 1995). For FAR estimates, this corresponded to an adjustment based on size of the risk set at the start of the analysis window (‘𝑡_𝑠_ ’) (Kalbfleisch and Prentice, 2011).

Simulations (**Fig. S2A**) and a lack of positive correlation between bias and sensitivity differences between blocks or sessions (see **Figs. 3** **and S3**) support that our approach to SDT successfully encapsulates and orthogonalizes d’ and c.

### Accumulation-to-bound modeling

We fitted the choice and reaction time data, pooled from all animals, with accumulation-to-bound models of the decision variable. Modles (Fig. 6A**-B**) were fitted based on continuous maximum likelihood using the Python-package PyDDM (Shinn et al., 2020). The combination of model parameters determines the fraction of correct responses and their associated RT distributions (Fig. 6C**-E**). We employed a single accumulator model that describes the accumulation of noisy sensory evidence toward a single choice boundary for a go-response.

In the model, the decision dynamics were governed by leak, drift bias and gaussian noise during ‘noise’ (tone cloud) stimuli, and additionally by the drift rate and signal neglect probability during the ‘signal’ (temporal coherence) stimuli, based on the following equation:

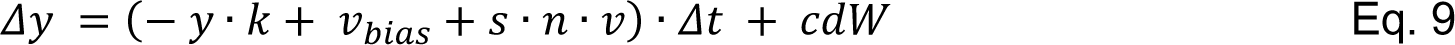

where 𝑦 is the decision variable (black example trace in Fig. 6A, right), 𝑘 is the leak and controls the effective time constant (1/𝑘) for which accumulated evidence decays, 𝑣_𝑏𝑖𝑎𝑠_ is an evidence-independent constant that is added as a drift in the diffusion process, 𝑠 is the stimulus category (0 during ‘noise’; 1 during ‘signal’), 𝑛, equal to (1 − 𝑝), is a Bernoulli variable (‘0’ or ‘1’), determined with probability 𝑝 as the fraction of signal presentations on which the relevant sensory evidence 𝑠 was neglected (not accumulated), 𝑣 is the drift rate and controls the overall efficiency of accumulation of relevant evidence (coherence), and 𝑐𝑑𝑊 is Gaussian distributed white noise with mean 0 and variance 𝑐^2^𝛥𝑡. Evidence accumulation terminated at the bound height (go response) or at the end of the trial (no-go response), whichever came first. The starting point of evidence accumulation was fixed to ‘0’.

Changes in bound height, leak and drift predict similar enough changes in the fraction of go-responses and fits that it proved to be unstable when letting all of these parameters vary freely with task utility. Therefore, we initially compared three different models: in model 0 all parameters could vary with task utility except bound height, in model 1 all parameters could vary except leak, and in model 2 all parameters could vary except drift bias. Model 0 produced the best fits, both quantitatively (Fig. 6B) as well as qualitatively (compare Fig. 6C**-E** to **Fig S6A-D**).

We fitted two additional alternative models to verify that the signal neglect probability was an essential parameter. Model 3 did not include signal neglect probability at all, and in model 4 signal neglect probability was fitted but could not vary with task utility. Both alternative models produced worse fits, both quantitatively (Fig. 6B) and qualitatively (**Fig S6E-H**).

We fitted two additional alternative models to verify that drift bias was an essential parameter. Model 5 did not include drift bias at all, and in model 6 drift bias was fitted but could not vary with task utility. Both alternative models produced worse fits quantitatively (Fig. 6B) and qualitatively (**Fig S6I-L**).

For the winning model (model ‘0’ we let all parameters except bound height vary with block number (Fig. 7A**-E**). In a separate analysis, all parameters except bound height could vary with the different arousal states and for the low-reward and high-reward blocks (**Fig. S7F-J)**. For simplicity of illustration and to pinpoint the effects of arousal on decision-making that are independent of task utility we then averaged the fits across task utility (Fig. 7F**-J**).

### Time course of learning

To characterize animal’s learning, we fitted the following function:

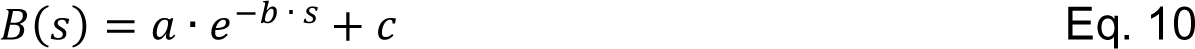

where 𝐵 is a behavioral metric of interest, 𝑠 is session number with respect to start of phase 3, and 𝑎, 𝑏 and 𝑐 the free parameters of the fit.

*Time course of behavioral adjustments.* To calculate the speed of within-block behavioral adjustments to changes in task utility, we fitted the following function:

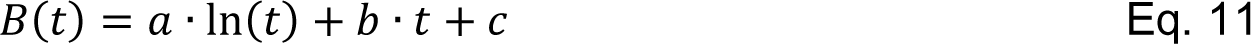

where 𝐵 is a behavioral metric of interest, 𝑡 is trial number since first correct response (hit) in a block, and 𝑎, 𝑏 and 𝑐 the free parameters of the fit. We then calculated the difference between the maximum and minimum of the fitted function and calculated the trial number for which 95% of this difference was reached. For bias, sensitivity, RT and reward probability, this occurred on average at 15 trials after the first correct response in a low-reward block (**Fig. 2B,E**,**H,K**). Therefore, when pooling data across trials within a block, we always excluded the first 14 trials after the first hit in each block (in both high-reward and low-reward blocks). We verified that our conclusions were not affected by specifics of this trial-selection procedure (**Fig. S2H-K**).

### Simulation of optimal signal-independent response rate

To characterize the theoretical relationship between an overall (signal-independent) Poisson response rate and our behavioral metrics we generated a simulated data set (**Fig. S2A**). Specifically, we generated synthetic trials that matched the statistics of the empirical trials (noise duration was drawn from an exponential distribution with mean = 5 s; truncated at 11 s, the signal duration was 3 s). We then systematically varied the overall response rate by drawing random response times from exponential distributions with various means (1 / rate) and randomly assigned those response times to the synthetic trials. We varied the overall response rate from 0.05 to 1 responses/s, in 20 evenly spaced steps. For each response rate, the decision agent performed 500 thousand simulated trials. For each iteration we then calculated the resulting bias, sensitivity, RT, and reward probability.

### Logistic regression

We split each trial into a noise and signal epoch (for false alarm trials only a noise epoch), and then fitted three different logistic regression models. First:

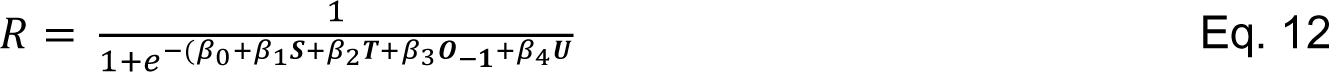

where ***R*** was a binary vector describing whether the animal responded (licked) during a given noise or signal epoch (0, no response; 1, response); ***S*** was a binary vector describing signal identity (0, noise; 1, signal); ***T*** was a vector describing trial number; ***O_-1_***was a binary vector describing whether the previous trial was rewarded (0, no previous reward; 1, previous reward); ***U*** was binary a vector describing the current task utility (0, low; 1, high); the *β*’s were the parameters (coefficients) of the fit. We next extended the model as follows:

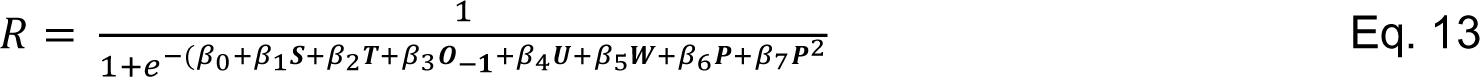

where ***W*** was binary a vector describing whether the animal walked (0, still; 1, walk); ***P*** was a vector describing the pre-trial pupil size (note that we included both a linear and quadratic term). We finally extended the model as follows:

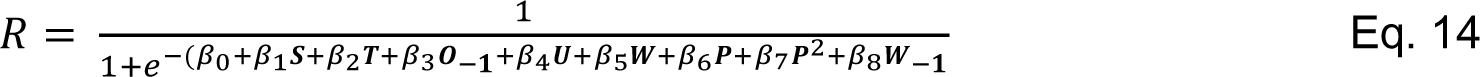

where ***W_-1_*** was binary a vector describing whether the animal walked on the previous trial (0, still; 1, walk).

### Mediation analysis

We used mediation analysis (Baron & Kenny, 1986) to characterize the interaction between task utility, arousal, and behavioral performance (Fig. 5I,**J**). We fitted the following linear regression models based on standard mediation path analysis:

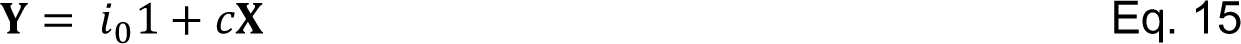

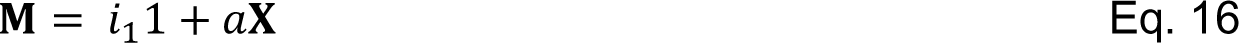

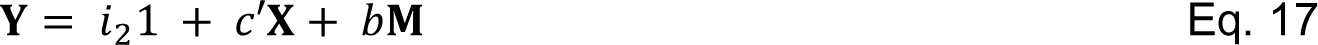

where ***Y*** was a vector of the block-wise behavioral metric (e.g., sensitivity), ***X*** was binary a vector describing the block-wise task utility (0, low; 1, high), **M** was a vector of block-wise distance with respect to the optimal measures (see *Analysis of pupil data*), and 𝑐, 𝑐′, 𝑎, 𝑏, 𝑖_0_, 𝑖_1_ and 𝑖_2_ 𝑤ere the free parameters of the fit. The parameters were fit using freely available Python software (Vallat, 2018).

### Analysis of pupil data

All analyses were performed using custom-made Python scripts, unless stated otherwise.

*Preprocessing.* We measured pupil size and exposed eye area from the videos of the animal’s eye using DeepLabCut (Mathis et al., 2018; Mridha et al., 2021). In approximately 1000 training frames randomly sampled across all sessions, we manually identified 8 points spaced approximately evenly along the edge of the pupil, and 8 points along the edge of the eyelids. The network (resnet 110) was trained with default parameters. To increase the network’s speed and accuracy when labeling (unseen) frames of all videos, we specified video-wise cropping values in the DeepLabCut configuration file that corresponded to a square around the eye. The pupil size (or exposed eye area) was computed as the area of an ellipse fitted to the detected pupil (exposed eye) points. If two or more points were labeled with a likelihood smaller than 0.1 (e.g., during blinks), we did not fit an ellipse, but flagged the frame as missing data. We then applied the following signal processing to the pupil (exposed eye) time series of each measurement session: (i) resampling to 10 Hz; (ii) blinks were detected by a custom algorithm that marked outliers in the z-scored temporal derivative of the pupil size time series; (iii) linear interpolation of missing or poor data due to blinks (interpolation time window, from 150 ms before until 150 ms after missing data); (iv) low-pass filtering (third-order Butterworth, cut-off: 3 Hz); and (v) conversion to percentage of the 99.9 percentile of the time series. See (McGinley, David, et al., 2015; Mridha et al., 2021) for additional details.

*Quantification of pre-trial pupil size and distance w.r.t optimal.* We quantified pre-trial pupil size as the mean pupil size during the 0.25 s before trial onset. Pre-trial pupil size was highest after previous hits (**Fig. S4A**), likely because the phasic lick-related pupil response did not have enough time to return to baseline. Pre-trial pupil size also generally increased with time-on-task (**Fig. S5G**). We thus removed (via linear regression) components explained by previous outcome (reward vs. no reward) and trial number. We obtained qualitatively similar results without doing so (**Fig. S4K**, **Fig. S5G-M**, and **Fig. S7K-T**). To capture how close the animal’s arousal state on each trial was to the optimal level, we computed the absolute difference between each pre-trial’s pupil size and the optimal size. Here, optimal size (which was found to be 27.5% of max) was defined as the pre-trial baseline pupil size for which reward probability was maximal (green vertical line in Fig. 4D).

*Utility context-based pupil resampling simulations.* Per animal and block type (high-reward and low-reward) we counted the number of trials in each arousal-defined bin (same as bins as in Fig. 5J); we then used these counts to compute the arousal-predicted behavioral performance (e.g., sensitivity), using the previously observed relationship between arousal states and behavioral performance (irrespective of task utility; Fig. 4A**-D**); per animal, we then computed the difference between the arousal-predicted behavioral performance in the high-reward and low-reward blocks. Finally, we divided the average (across animals) arousal-predicted difference by the average (across animals) total actually observed difference (Fig. 2). To estimate uncertainty, we bootstrapped trials within animals and blocks (5K bootstraps). We then computed the fraction between the purely arousal-predicted change in behavior and the total change in behavior after changes in task utility (see Fig. 5K).

### Analysis of walking data

The instantaneous walking speed data was resampled to 10 Hz. We quantified pre-trial walking speed as the mean walking velocity during the 2 s before trial onset. We defined walking probability as the fraction of trials for which the absolute walking speed exceeded 1.25 cm/s (**Fig. S4B**).

### Statistical comparisons

We used a 3 × 2 repeated measures ANOVA to test for the main effects of task utility and time-on-task (block number of a given reward magnitude), and their interaction (Fig. 2C,**F**,**I,L** and Fig. 5C,**F**). We used the non-parametric Wilcoxon signed rank test to test coefficients against 0 (Fig. 4B,**C** and Fig. 5H,**J**).

We used sequential polynomial regression analysis (Draper & Smith, 1998), to quantify whether the relationships between pre-trial pupil size and behavioral measures were better described by a 1^st^ order (linear) or 2^nd^ order model (Fig. 4A**-D** and Fig. 7F**-J**):

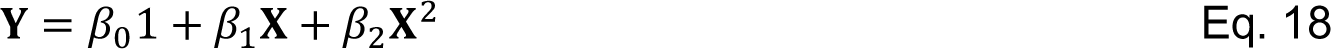

where **Y** was a vector of the dependent variable (e.g., bin-wise sensitivity), **X** was a vector of the independent variable (e.g. bin-wise pre-trial pupil size), and *β* as polynomial coefficients. To assess the amount of variance that each predictor accounted for independently, we orthogonalized the regressors prior to model fitting using QR-decomposition. Starting with the zero-order (constant) model and based on F-statistics (Draper & Smith, 1998), we tested whether incrementally adding higher-order predictors improves the model significantly (explains significantly more variance). We tested 1^st^ up to 5^th^ order models.

We used Bayesian information criterion (BIC) for model selection and verified whether the complexity of the different variants of the accumulation-to-bound model was justified to account for the data (Fig. 6B). A difference in BIC of 10 is generally taken as a threshold for considering one model a sufficiently better fit than another (Spiegelhalter et al., 2002). We directly compared bootstrapped distributions of the model parameter estimates to test for the main effects of task utility and time-on-task (Fig. 7A**-E**) and for the effects pupil-linked and walk-related arousal (Fig. 7F**-J**).

All tests were performed two-tailed. All error bars are 68% bootstrapped confidence intervals of the mean, unless stated otherwise.

### Data availability

Data will be made publicly available upon publication.

### Code availability

Analysis scripts will be made publicly available upon publication.

## Acknowledgements

We thank Daeyeol Lee for help designing the behavioral paradigm, Anton Banta for technical assistance, Sarim Aleem for help with pupil size analysis, many Rice University undergraduate students for help with animal behavioral training, Max Shinn, Konstantinos Tsetsos and Peter Murphy for helpful discussions about the accumulation-to-bound modeling and Nelson Totah for helpful comments on an early draft of the manuscript. Funding: NIDCD R01 and R03.

## Author Contributions

Conceptualization: JWG, MJM

Funding Acquisition: MJM

Data acquisition software: WZ, HJ

Investigation: ZM, MH, YS, HR, SS, NK, MT, KJ

Data Curation: JWG, MH, YS, HR, SS, NK, MT, KJ

Formal analysis: JWG, MJM

Methodology: JWG, MJM

Writing—original draft: JWG, MJM

Writing—review and editing: JWG, ZM, MJM

Supervision: MJM

Project Administration: MJM

## Supplementary figures S1-S7

**Figure S1.**
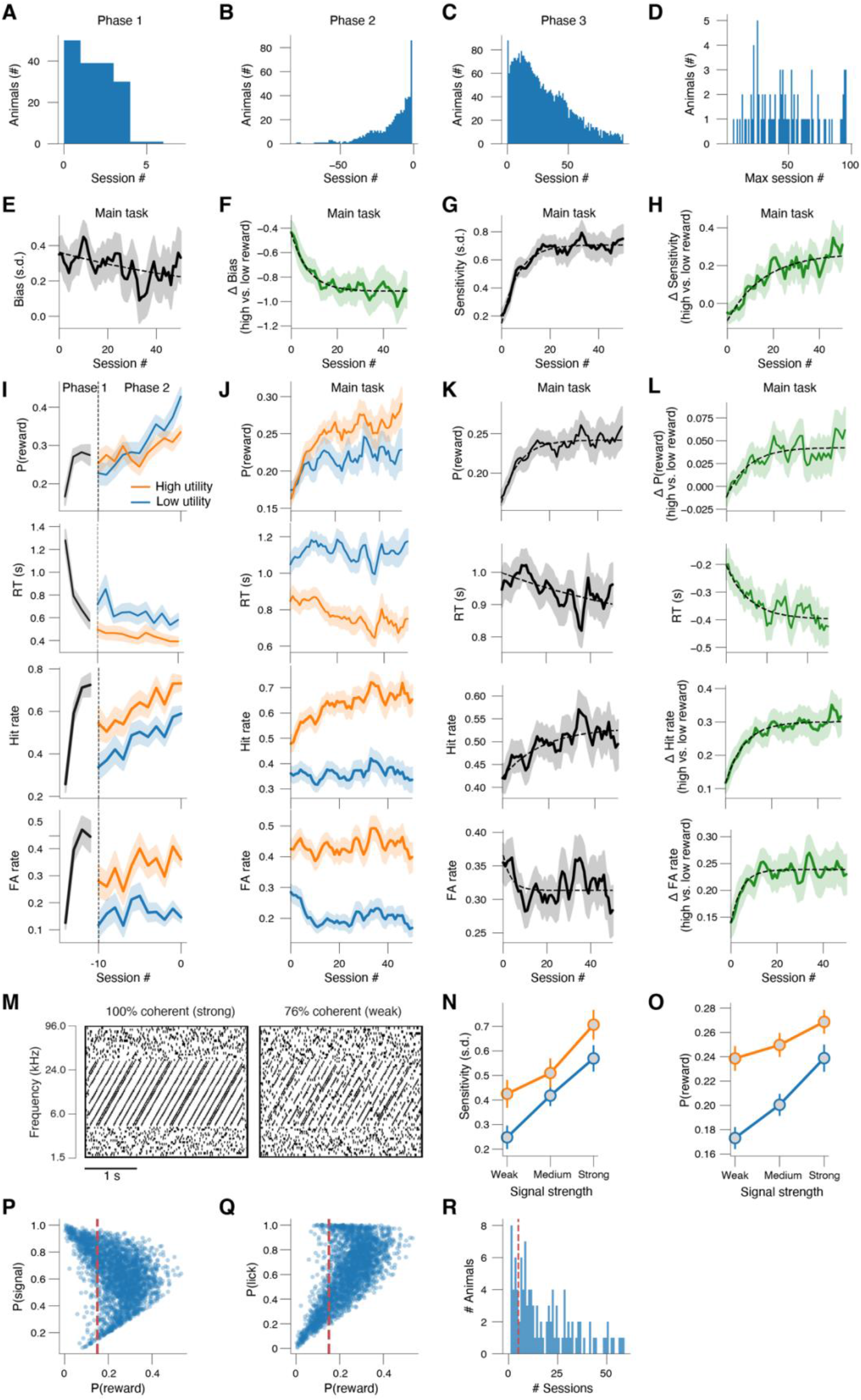
**(A)** Histogram of experimental session number in learning phase 1. **(B)** Histogram of experimental session number in learning phase 2 (with respect to last session number in phase 2). **(C)** Histogram of experimental session number in phase 3. **(D)** Histogram of the maximum session number per animal. **(E)** Bias across experimental sessions in learning phase 3, collapsed across reward context. **(F)** As E, but for the difference between high-reward and low-reward blocks. Dashed lines, exponential fit (Methods). **(G-H)** As E-F, but for sensitivity (Methods). **(I)** From top to bottom: reward probability, reaction time (RT), hit rate and false alarm rate (Methods) across experimental sessions in learning phases 1 and 2 (Methods); session numbers are with respect to the last session in phase 2. **(J)** As I, but for the main task (Methods). **(K)** As I, but collapsed across reward context. **(L)** As I, but for the difference between high-reward and low-reward blocks. Dashed lines, exponential fit (Methods). **(M)** Example spectrograms of strong (left) and weak (right) signals (Methods). **(N)** Sensitivity across three difficulty bins and separately for high-reward and low-reward blocks. Stats, 2-way repeated measures ANOVA (factors task utility [high vs. low] and signal strength (coherence) bin [weak, medium, strong]); main effect task utility: F_1,9_ = 4.8, p = 0.056; main effect signal strength: F_2,18_ = 29.7, p < 0.001; interaction effect: F_2,18_ = 1.7, p = 0.201. Error bars, 68% confidence interval across animals (N=10, n=142 sessions). **(O)** As N, but for reward probability. Main effect task utility: F_1,9_ = 17.8, p = 0.002; main effect signal strength: F_2,18_ = 8.0, p = 0.003; interaction effect: F_2,18_ = 2.0, p = 0.162. **(P)** Fraction of trials containing a signal plotted against reward probability. Every data point is a unique session. We excluded 455 sessions with a reward probability smaller than 0.15, thereby excluding 4 animals. **(Q)** As P, but for fraction of trials on which the animal licked (responded) on the y-axis. **(R)** Histogram of number of sessions per animal (after excluding sessions in panel Q,R). We excluded 22 additional animals with fewer than 5 remaining sessions.

**Figure S2.**
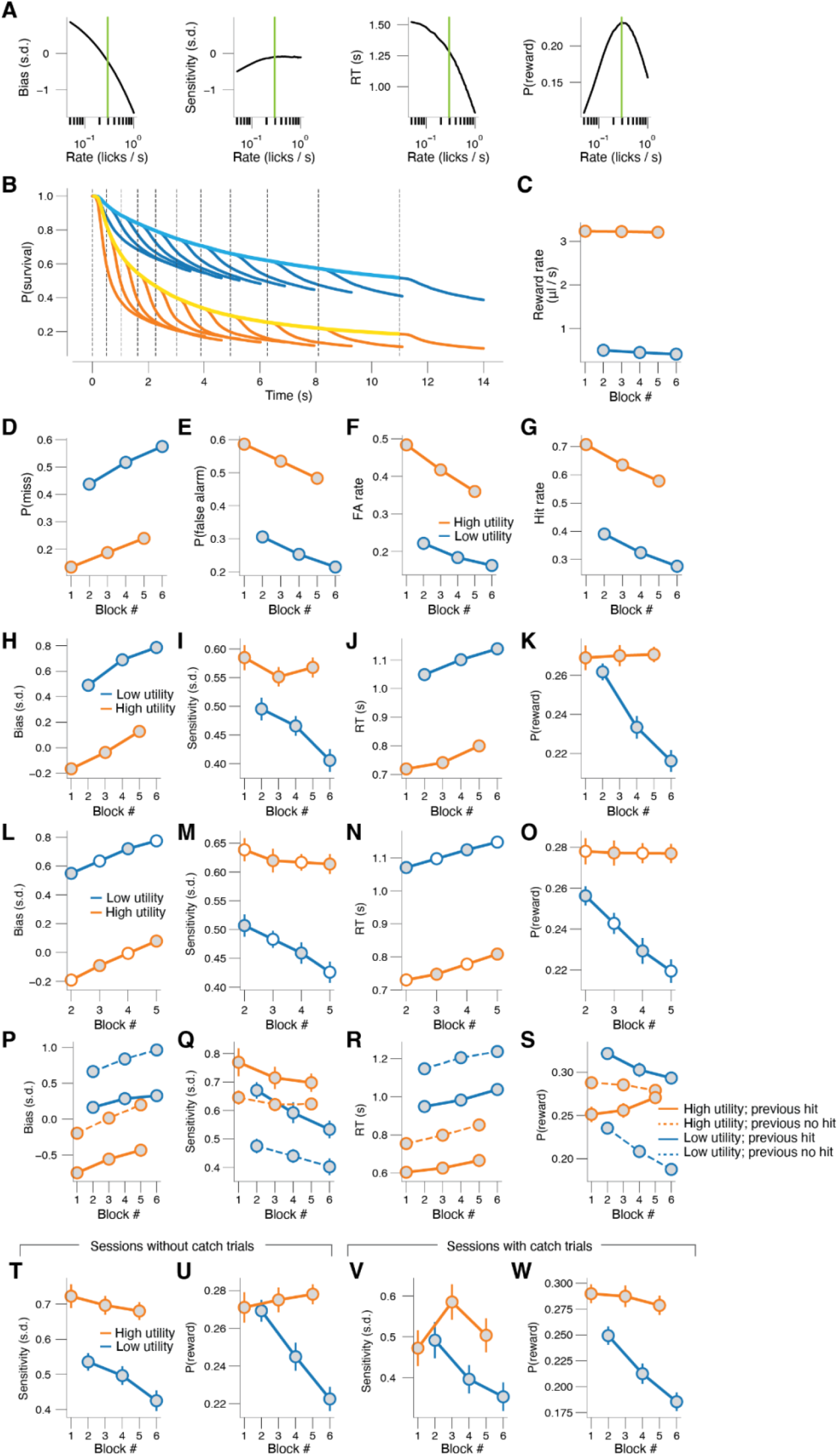
**(A)** From left to right: simulation of bias, sensitivity, RT and reward probability as a function of Poisson lick rate (Methods). Vertical green line indicates the optimal lick rate (maximum reward probability) when lick rate during noise and signal are the same. Note: the optimal average rate may differ in conditions when the signal is discriminated from the noise. **(B)** Probability of survival (Kaplan-Meier estimator), separately for low-reward and high-reward blocks, separately for noise and signal epochs, and separately for different noise durations. **(C)** Reward rate (Methods) collapsed across trials within each block. Stats, 2-way repeated measures ANOVA (factors task utility [high vs. low] and time-on-task [1, 2, 3]); main effect task utility: F_1,87_ = 1519.3, p < 0.001; main effect time-on-task: F_2,174_ = 1.9, p = 0.155; interaction effect: F_2,174_ = 0.7, p = 0.519. **(D)** As C but for miss probability (Methods). Main effect task utility: F_1,87_ = 985.2, p < 0.001; main effect time-on-task: F_2,174_ = 177.1, p < 0.001; interaction effect: F_2,174_ = 1.5, p = 0.228. **(E)** As C but for false alarm probability (Methods). Main effect task utility: F_1,87_ = 714.5, p < 0.001; main effect time-on-task: F_2,174_ = 188.2, p < 0.001; interaction effect: F_2,174_ = 4.2, p = 0.016. **(F)** As C but for false alarm rate (Methods). Main effect task utility: F_1,87_ = 439.0, p < 0.001; main effect time-on-task: F_2,174_ = 86.6, p < 0.001; interaction effect: F_2,174_ = 17.9, p < 0.001. **(G)** As C but for hit rate (Methods). Main effect task utility: F_1,87_ = 538.1, p < 0.001; main effect time-on-task: F_2,174_ = 139.4, p < 0.001; interaction effect: F_2,174_ = 0.7, p = 0.512. **(H-K)** As C and main Fig. 2C,F,I,L, but when using all trials within a block (Methods). Main effects task utility are as follows: Panel H: F_1,87_ = 747.0, p < 0.001. Panel I: F_1, 87_ = 17.7, p < 0.001. Panel J: F_1, 87_ = 612.9, p < 0.001. Panel K: F_1, 87_ = 12.5, p = 0.001. **(L-O)** As C and main Fig. 2C,F,I,L, but when controlling for time-on-task. The white circles are the average of the two adjacent blocks of the same task utility. Main effects task utility are as follows: Panel L: F_1,87_ = 498.5, p < 0.001. Panel M: F_1, 87_ = 26.8, p < 0.001. Panel N: F_1, 87_ = 365.1, p < 0.001. Panel O: F_1, 87_ = 15.3, p < 0.001. **(P-S)** As C and main Fig. 2C,F,I,L, but when additionally stratifying on previous hit (Methods). Main effects previous hit are as follows: Panel P: F_1,87_ = 733.7, p < 0.001. Panel Q: F_1, 87_ = 34.0, p < 0.001. Panel R: F_1, 87_ = 184.2, p < 0.001. Panel S: F_1, 87_ = 55.1, p < 0.001. Main effects task utility are as follows: Panel P: F_1,87_ = 770.2, p < 0.001. Panel Q: F_1, 87_ = 21.9, p < 0.001. Panel R: F_1, 87_ = 667.0, p < 0.001. Panel S: F_1, 87_ = 1.8, p = 0.179. **(T**,**U)** As main Fig. 2F,L, but only for 82.8% of sessions without catch trials (Methods). Main effects task utility are as follows: Panel S: F_1,69_ = 31.9, p < 0.001. Panel T: F_1, 69_ = 6.2, p = 0.015. **(V,W)** As main Fig. 2F,L, but only for 17.2% of sessions with catch trials (Methods). Main effects task utility are as follows: Panel V: F_1,43_ = 6.6, p = 0.013. Panel W: F_1, 43_ = 29.0, p < 0.001. All panels: shading or error bars, 68% confidence interval across animals (N=88; n=1983 sessions).

**Figure S3.**
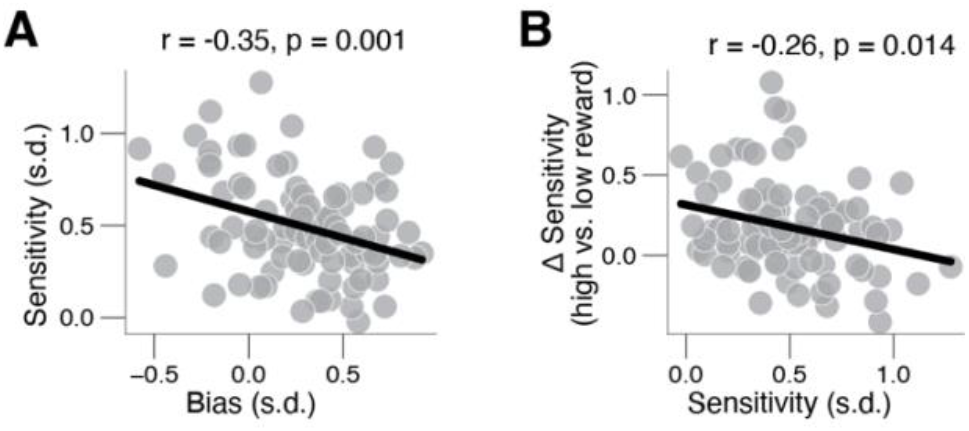
**(A)** Overall sensitivity (collapsed across task utility) plotted against overall bias (collapsed across task utility). Every data point is a unique session. **(B)** Change in sensitivity between reward blocks plotted against overall sensitivity (collapsed across task utility). Every data point is a unique session.

**Figure S4.**
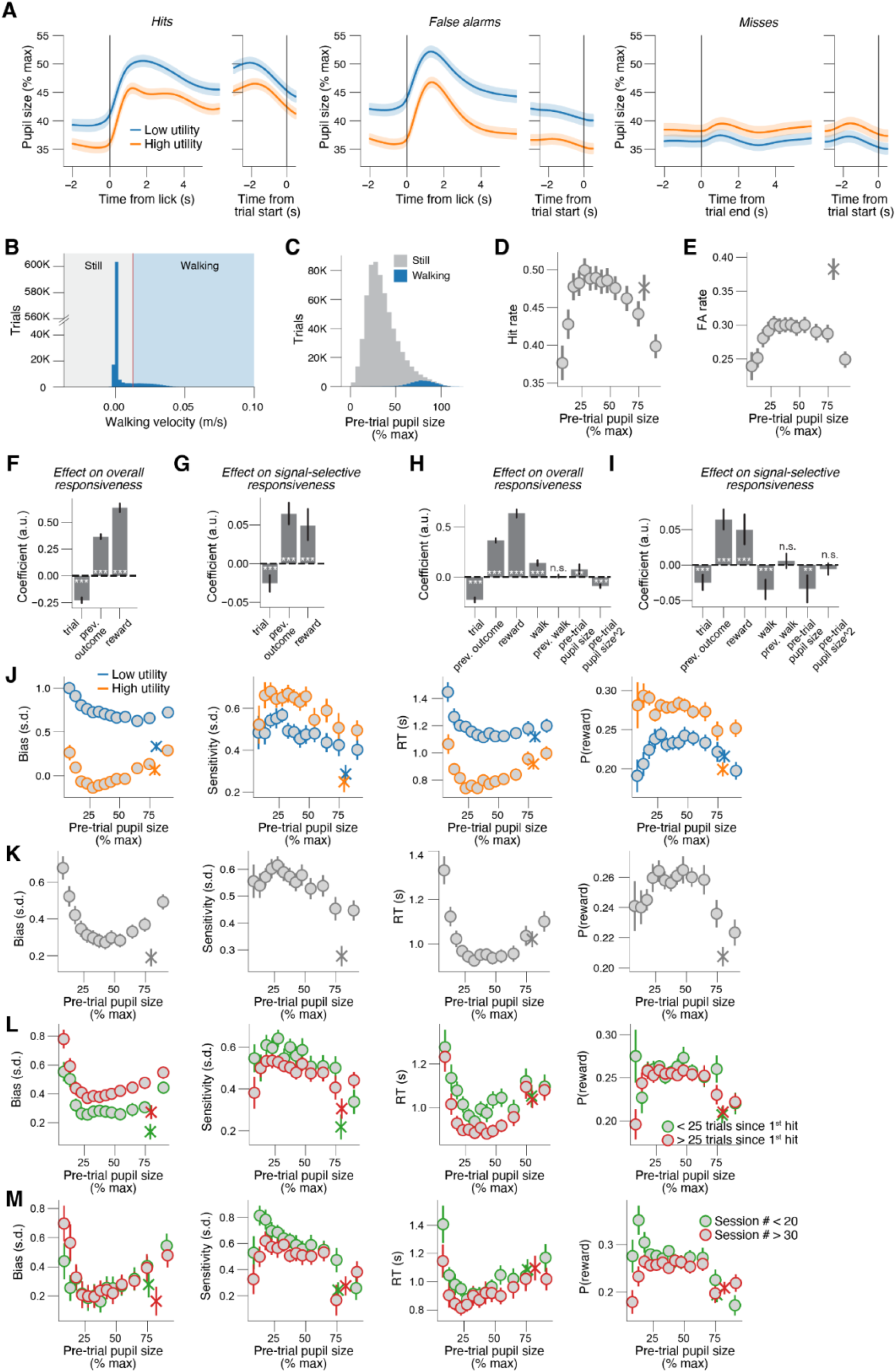
**(A)** Left: Pupil size across time on hit trials, separately for high-reward and low-reward blocks. Left, locked to licks; right, locked to next trial’s onset. Middle, as left, but on false alarm trials. Right: As left, but for miss trials. Left, locked to trial offset (end of 3-s signal sound). **(B)** Histogram of pre-trial walking velocity (across all animals and experimental sessions; Methods). Red line, cutoff for defining walking. **(C)** Histogram of pre-trial pupil size (across all animals and experimental sessions), separately for still and walking trials. **(D)** Relationship between pre-trial pupil size and hit rate (irrespective of task utility; Methods). A 1^st^ order (linear) fit was not superior to a constant fit (F_1,12_ = 0.1, p = 0.717) and a 2^nd^ order (quadratic) fit was superior to the 1^st^ order fit (F_1,12_ = 6.5, p = 0.026; sequential polynomial regression; Methods). Asterisk, walking trials (Methods). **(E)** As D, but for false alarm rate. 1^st^ order fit: F_1,12_ = 0.2, p = 0.690; 2^nd^ order fit: F_1,12_ = 7.9, p = 0.016. **(F)** Remaining fitted coefficients from multiple logistic regression model (Fig. 4E; Methods), capturing the effects of time-on-task, previous outcome, and utility on overall responsiveness (closely related to bias). Stats, Wilcoxon signed-rank test; **, p < 0.01; ***, p < 0.001. **(G)** As F, but for interaction effects between each factor and signal, capturing the effects of time-on-task, previous hit, and utility on signal-selective responsiveness (closely related to sensitivity). **(H,I)** As F,G, but for model with one additional predictor: previous walk. **(J) As** Fig. 4A-D, but separately for low-reward and high-reward blocks. **(K)** As Fig. 4A-D, but for pre-trial pupil size measures without having regressed out effects of time-on-task and previous hit (Methods). Stats (sequential polynomial regression; Methods) are as follows. Bias: 1^st^ order fit: F_1,12_ = 13.2, p = 0.003; 2^nd^ order fit: F_1,12_ = 8.0, p = 0.016. Sensitivity: 1^st^ order fit: F_1,12_ = 4.4, p = 0.058; 2^nd^ order fit: F_1,12_ = 3.0, p = 0.111. RT: 1^st^ order fit: F_1,12_ = 5.1, p = 0.043; 2^nd^ order fit: F_1,12_ = 7.5, p = 0.018. Reward probability: 1^st^ order fit: F_1,12_ = 1.3, p = 0.268; 2^nd^ order fit: F_1,12_ = 7.3, p = 0.019. **(L)** As Fig. 4A-D, but separately for fewer or more than 25 trials after the first hit in each block. **(M)** As Fig. 4A-D, but separately for the first 20 sessions of the main task, or for session numbers greater than 30. In this analysis, we only considered the first 50 sessions of 34 animals with at least 50 sessions worth of data in the main task. All panels: shading or error bars, 68% confidence interval across animals (N=88, n=1983 sessions).

**Figure S5.**
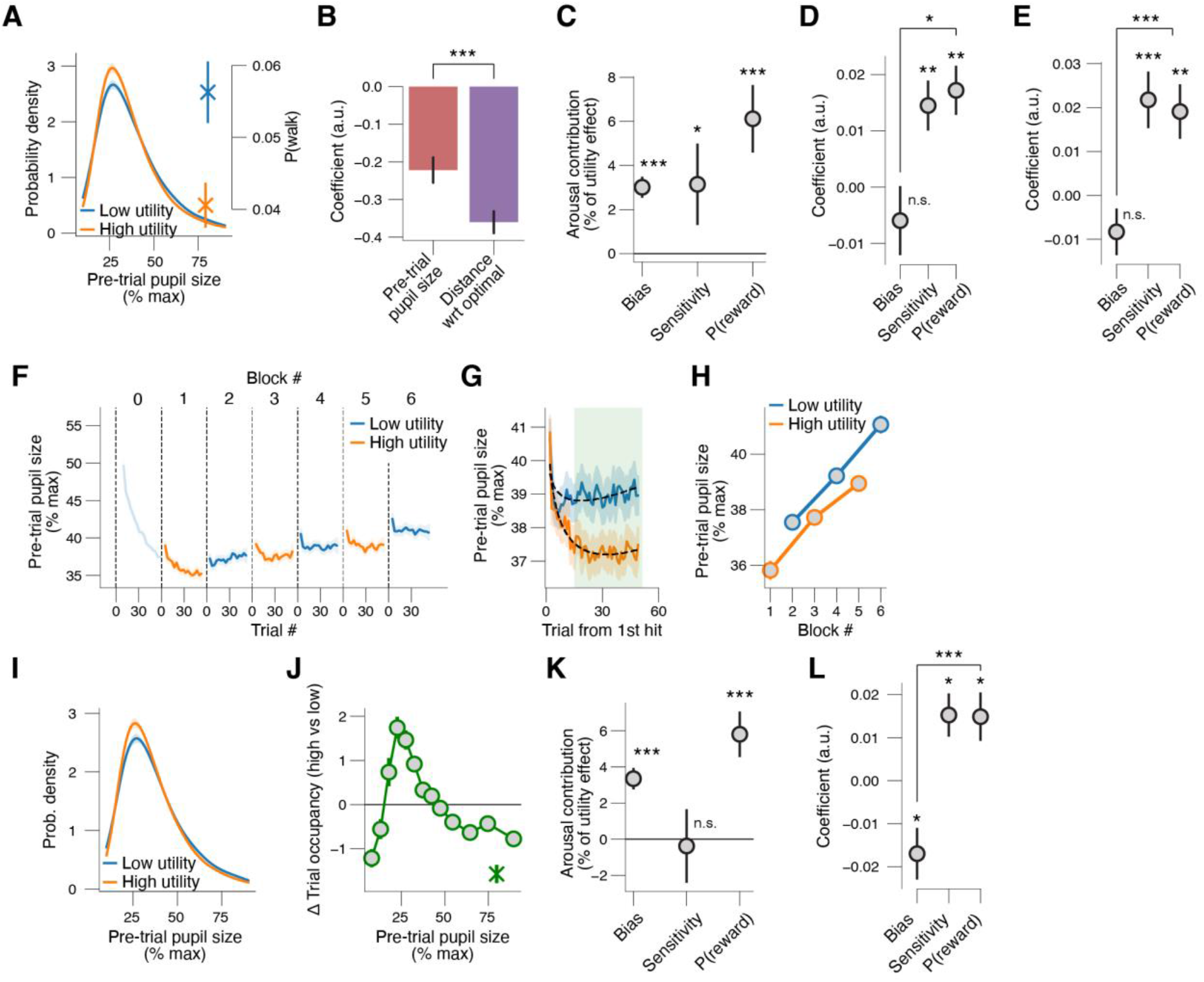
**(A)** Probability density function of pupil-linked arousal states, and walk probability, separately for low-reward and high-reward blocks. The difference between is reported in main Fig. 5G. **(B)** Coefficients from logistic regression of block-wise task utility [high vs. low] on pre-trial pupil size or on absolute distance from optimal pre-trial pupil size (Methods). **(C)** As main Fig. 5H, but without modeling walking trials in a separate state bin. **(D)** As main Fig. 5J, but only for trials mice did not walk. **(E)** As main Fig. 5J, but with walk probability as mediator (pupil-linked arousal not considered). **(F)** Pre-trial pupil size (without having regressed out time-on-task and previous hit; Methods) across low-reward and high-reward blocks in each experimental session, locked to first hit in block. Data from the first block of each session (low utility; termed block ‘0’) was excluded from all analyses, as mice spent this block becoming engaged in the task (see also Fig. 2A,D,G,J; Methods). **(G)** As G, but collapsed across blocks of same reward magnitude. The green shaded area indicates the trials used when pooling data across trials within a block (e.g. panel C). **(H)** As G, but collapsed across trials within a block. Stats, 2-way repeated measures ANOVA (factors task utility [high vs. low] and time-on-task [early, middle, late]); main effect task utility: F_1,87_ = 43.4, p < 0.001; main effect time-on-task: F_2,174_ = 37.1, p < 0.001; interaction effect: F_2,174_ = 3.3, p = 0.282. **(I)** As C, but for pre-trial pupil size without having regressed out time-on-task and previous hit (Methods). **(J)** Change in trial density after increases in task utility, separately for pupil-defined arousal states; asterisk, walking trials, without having regressed out time-on-task and previous hit (Methods). **(K)** As main Fig. 5H, but for pre-trial pupil size without having regressed out time-on-task and previous hit (Methods). **(L)** As main Fig. 5J, but for pre-trial pupil size without having regressed out time-on-task and previous hit (Methods). All panels: shading or error bars, 68% confidence interval across animals (N=88, n=1983 sessions). Panels C-E,K,L: stats, Wilcoxon signed-rank test; *, p < 0.05; **, p < 0.01***, p < 0.001.

**Figure S6.**
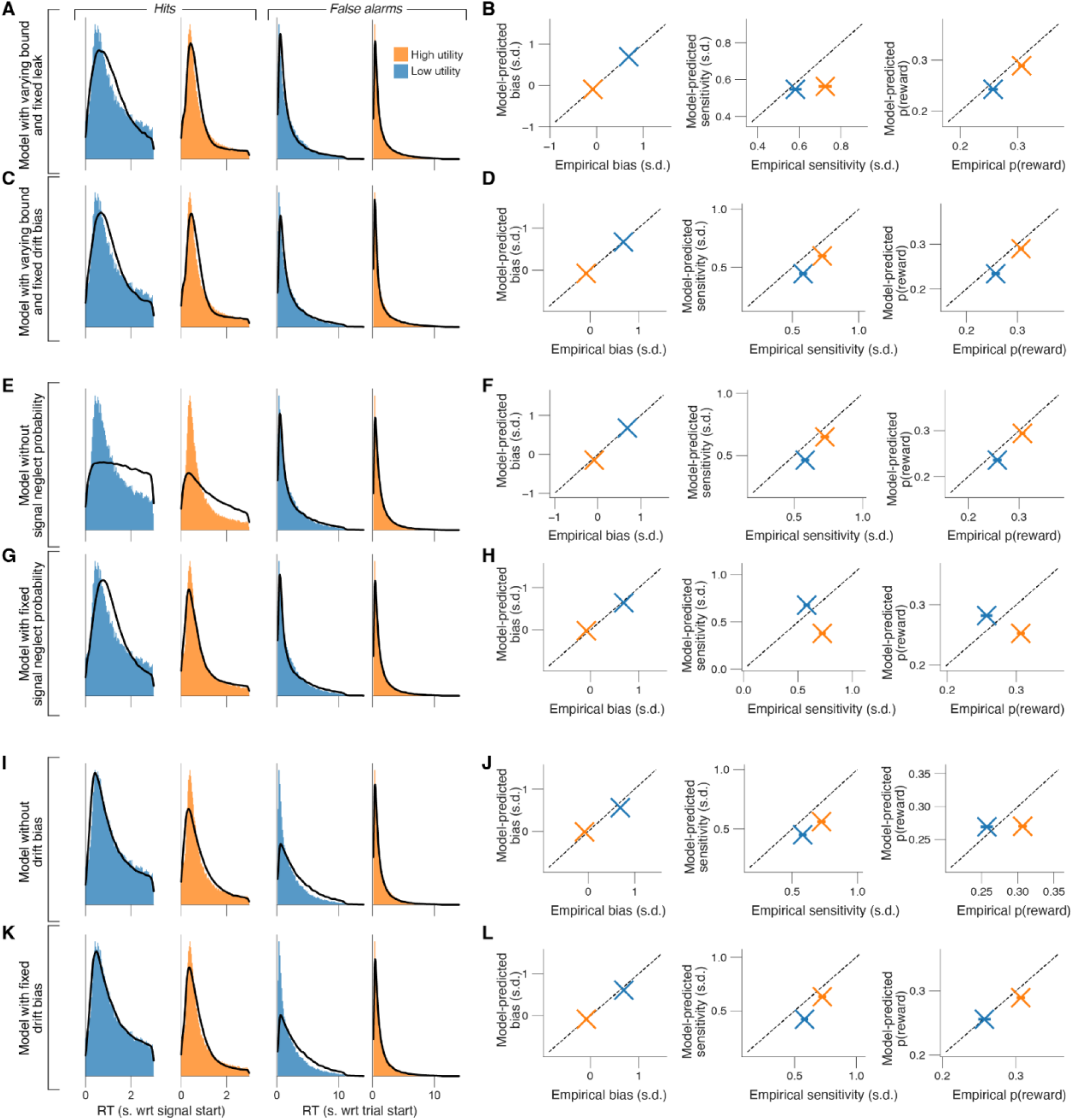
**(A)** RT distribution for correct responses (hits; left) and incorrect responses (false alarms; right) in the low-reward and high-reward blocks. Black line, fit of Model 1 (with varying bound height and fixed leak; Methods). **(B)** Model-predicted bias (left), sensitivity (middle) and reward probability (right) in the low-reward and high-reward blocks plotted against the empirical estimates. Dashed line, identity line. **(C,D)** As A,B, but for Model 2 (with varying bound height and fixed drift bias; Methods). **(E,F)** As A,B, but for Model 3 (without signal neglect probability; Methods). **(G,H)** As A,B, but for Model 4 (with fixed signal neglect probability; Methods). (I,J) As A,B, but for Model 5 (without drift bias; Methods). **(K,L)** As A, B, but for Model 6 (with fixed drift bias; Methods). All panels: pooled data across animals (N=88) and sessions (n=1983).

**Figure S7.**
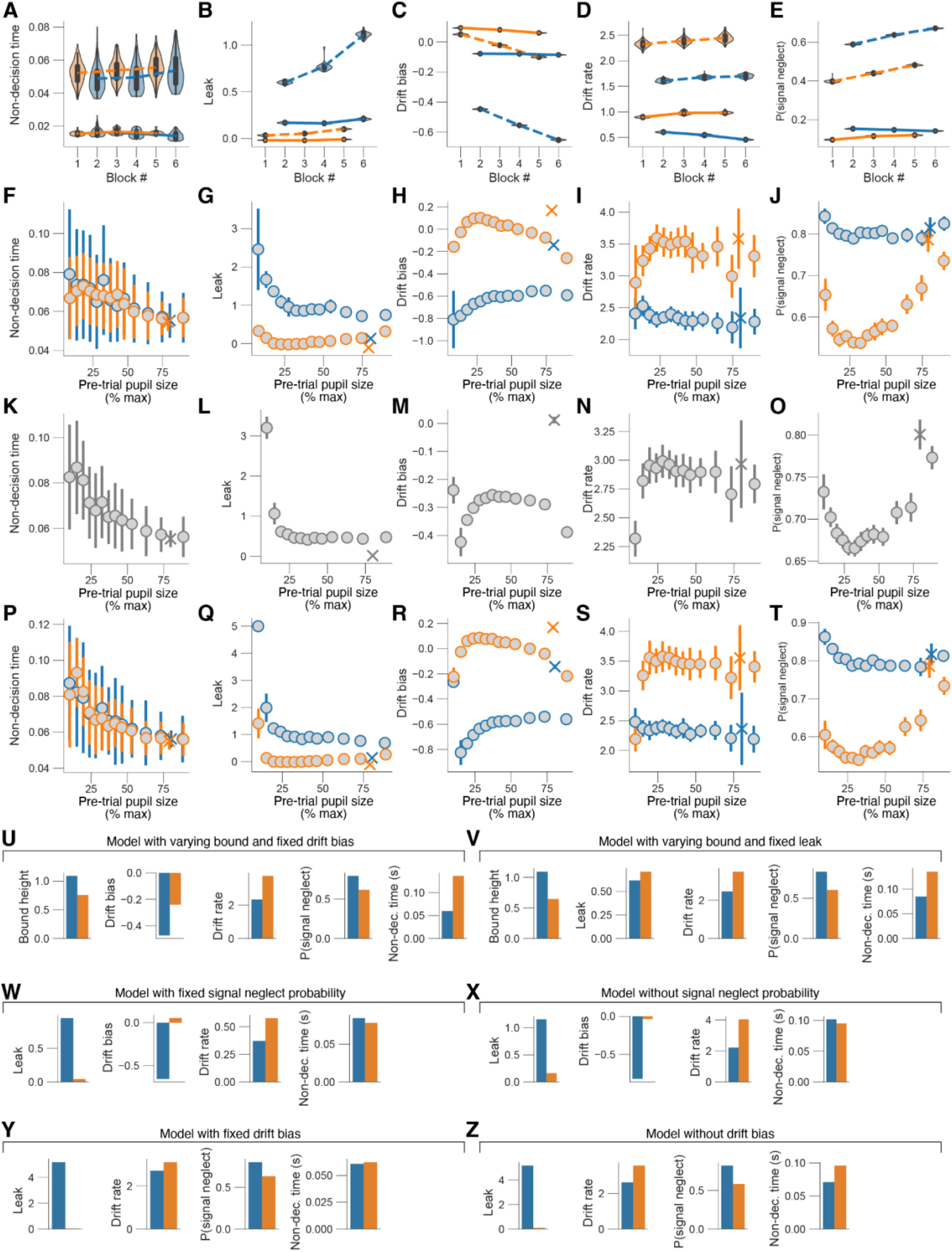
**(A)** Fitted non-decision time estimates (kernel density estimate of 100bootstrapped replicates) separately per block number. Main effect task utility (fraction of bootstrapped parameter estimates in the low-reward blocks higher than in the high-reward blocks): p = 0.32. Main effect time-on-task (fraction of bootstrapped parameter estimates in the first two blocks higher than in the last two blocks): p = 0.41. (B) As A, but for leak. Main effect task utility: p < 0.01. Main effect time-on-task: p < 0.01 (C) As A, but for drift bias. Main effect task utility: p < 0.01. Main effect time-on-task: p < 0.01. (D) As A, but for drift rate. Main effect task utility: p < 0.01. Main effect time-on-task: p = 0.46. (E) As A, but for signal neglect probability. Main effect task utility: p < 0.01. Main effect time-on-task: p < 0.01. (F-J) As main Fig. 7F-J, but separately for low-reward and high-reward blocks. (K) Fitted non-decision estimates (100 bootstrapped replicates) separately per arousal state (same pupil size defined bins as in Fig. 4A-D, but without having regressed out effects of time-on-task and previous hit; irrespective of task utility; Methods). A 1^st^ order (linear) fit was superior to a constant fit (F_1,12_ = 118.8, p < 0.001) and a 2^nd^ order (quadratic) fit was not superior to the 1^st^ order fit (F_1,12_ = 2.3, p = 0.154; sequential polynomial regression; Methods). Asterisk, walking trials (Methods). (L) As K, but for leak. 1^st^ order fit: F_1,12_ = 5.9, p = 0.032; 2^nd^ order fit: F_1,12_ = 3.3, p = 0.093. (M) As K, but for drift bias. 1^st^ order fit: F_1,12_ ∼ 0.0, p = 0.870; 2^nd^ order fit: F_1,12_ = 1.3, p = 0.269. (N) As K, but for drift rate. 1^st^ order fit: F_1,12_ = 0.6, p = 0.469; 2^nd^ order fit: F_1,12_ = 3.8, p = 0.076. (O) As K, but for signal neglect probability. 1^st^ order fit: F_1,12_ = 1.2, p = 0.290; 2^nd^ order fit: F_1,12_ = 7.9, p = 0.016. (P-T) As 7F-J, but separately for low-reward and high-reward blocks. (U-Z) Parameter estimates from alternative models (Methods), separately for low-reward and high-reward blocks. All panels: pooled data across animals (N=88) and sessions (n=1983).

